# Human Gut Bacteria Convert Endogenous Steroids into Host Cortisol Shuttle Inhibitors

**DOI:** 10.64898/2026.08.21.746344

**Authors:** Seraina O. Moser, Jasmine T. Walsh, Philipp M. Späne, Taina Härkönen, Marianne Lalli, Ines Zalosnik, Denise V. Winter, Junehee Park, Marina Magicheva-Gupta, Megan D. McCurry, David J. Morris, Ye-Ji Bang, Jun R. Huh, Perry A. Labelle, Andrew T. Chan, David A. Drew, Mingyang Song, James D. Lewis, Gary D. Wu, Jordan E. Bisanz, Tommi Vatanen, Rianne Elaine van Diest, Mikael Knip, Alex Odermatt, A. Sloan Devlin

## Abstract

11β-hydroxysteroid dehydrogenase 2 (HSD11B2) protects the mineralocorticoid receptor from glucocorticoid overstimulation by inactivating cortisol. HSD11B2 inhibition can drive receptor overactivation and contribute to hypertension; however, endogenous inhibitors remain poorly defined. Glycyrrhetinic acid-like factors (GALFs) are steroid-like metabolites that inhibit HSD11B2. Given the capacity of gut bacteria to metabolize host steroids, we hypothesized that the gut microbiome could generate GALF-like inhibitors. Here, we identify two 11-oxygenated steroid metabolites that potently inhibit human HSD11B2 in colonic cells and organoids, enabling cortisol-dependent mineralocorticoid receptor activation. We identify gut bacteria and enzymes that produce these compounds and show that their levels are markedly reduced in antibiotic-treated humans. One metabolite is elevated during pregnancy, and both are higher in individuals with stage 2 hypertension-range blood pressure. These findings reveal a bacterial route to altered host cortisol signaling and suggest a potential link between microbiome-derived GALFs and blood pressure regulation.

## INTRODUCTION

Hypertension affects nearly half of US adults and increases the risk of cardiovascular disease, kidney disease, and diabetes.^1,2^ Preeclampsia, a pregnancy-specific hypertensive disorder, is a major cause of maternal and fetal morbidity and mortality worldwide.^3,4^ A growing number of studies implicate the gut microbiome in blood pressure regulation. For example, antibiotic treatment lowers blood pressure in hypertensive rats,^5–7^ and germ-free mice receiving fecal microbiota transplants (FMTs) from hypertensive but not normotensive donors develop elevated blood pressure.^8^ Likewise, microbiota from preeclampsia patients induces elevated blood pressure and preeclampsia-like phenotypes in antibiotic-treated pregnant mice.^9^ In humans hypertension is associated with gut dysbiosis, including enrichment of *Eggerthellaceae*.^8,10^ Several microbiome-derived metabolites, including trimethylamine N-oxide (TMAO) and short-chain fatty acids (SCFAs), have been implicated in blood pressure regulation.^11–13^ However, the molecular mechanisms by which the gut microbiome contributes to hypertension remain poorly defined.

Mineralocorticoid receptor (MR) signaling regulates sodium and potassium balance and fluid homeostasis, making it a potential target for gut microbiome-derived metabolites that influence blood pressure.^14,15^ MR is highly expressed in sodium-transporting epithelial cells of the kidney and distal colon.^16–23^ Aldosterone, the primary mineralocorticoid, is released in response to low blood pressure and activates the MR to promote sodium reabsorption and potassium secretion.^14,15^ This response is mediated in part through upregulation of basolateral Na⁺/K⁺- ATPase and increased activity of epithelial sodium channels (ENaC), resulting in sodium retention, fluid expansion, and restoration of blood pressure.^24–27^ The abundant glucocorticoid cortisol also binds the MR with similar affinity to aldosterone, and circulating cortisol concentrations are ∼100-1000-fold higher than aldosterone under physiological conditions.^16,28^ MR ligand specificity is therefore maintained by HSD11B2, also known as SDR9C3, a key enzyme in the mammalian cortisol shuttle. This enzyme inactivates cortisol by converting it to cortisone, which has negligible capacity to bind the MR.^16,17,28–31^ HSD11B2 is co-expressed with the MR in sodium-transporting epithelial tissues, including the kidney and distal colon, where it protects the MR from glucocorticoid overstimulation.^17,32^ Thus, HSD11B2 serves as a critical gatekeeper of MR activation by controlling glucocorticoid access to the receptor.

Disruption of HSD11B2 activity leads to pathological MR activation and hypertension. Loss-of-function mutations cause apparent mineralocorticoid excess (AME), a juvenile hypertensive syndrome in which cortisol activates the MR, resulting in sodium retention and potassium secretion, while renin and aldosterone levels remain low.^33–39^ Pharmacological inhibition of HSD11B2 similarly produces an acquired mineralocorticoid excess phenotype and contributes to hypertension (**Fig. 1A**).^38, 40–44^ A well-characterized example is the licorice-derived triterpenoid 18β-glycyrrhetinic acid (GA), a potent HSD11B2 inhibitor (**Fig. 1A-C**).^28,40–44^ Prior work has proposed that a broader class of endogenous steroid and steroid-like metabolites, termed GALFs, may similarly inhibit HSD11B2 and promote MR activation.^40,45^ Notably, because HSD11B2 is highly expressed in colonic tissues, the gut microbiome presents a plausible source of GALF-like molecules.^32,40,46^

**Figure 1.**
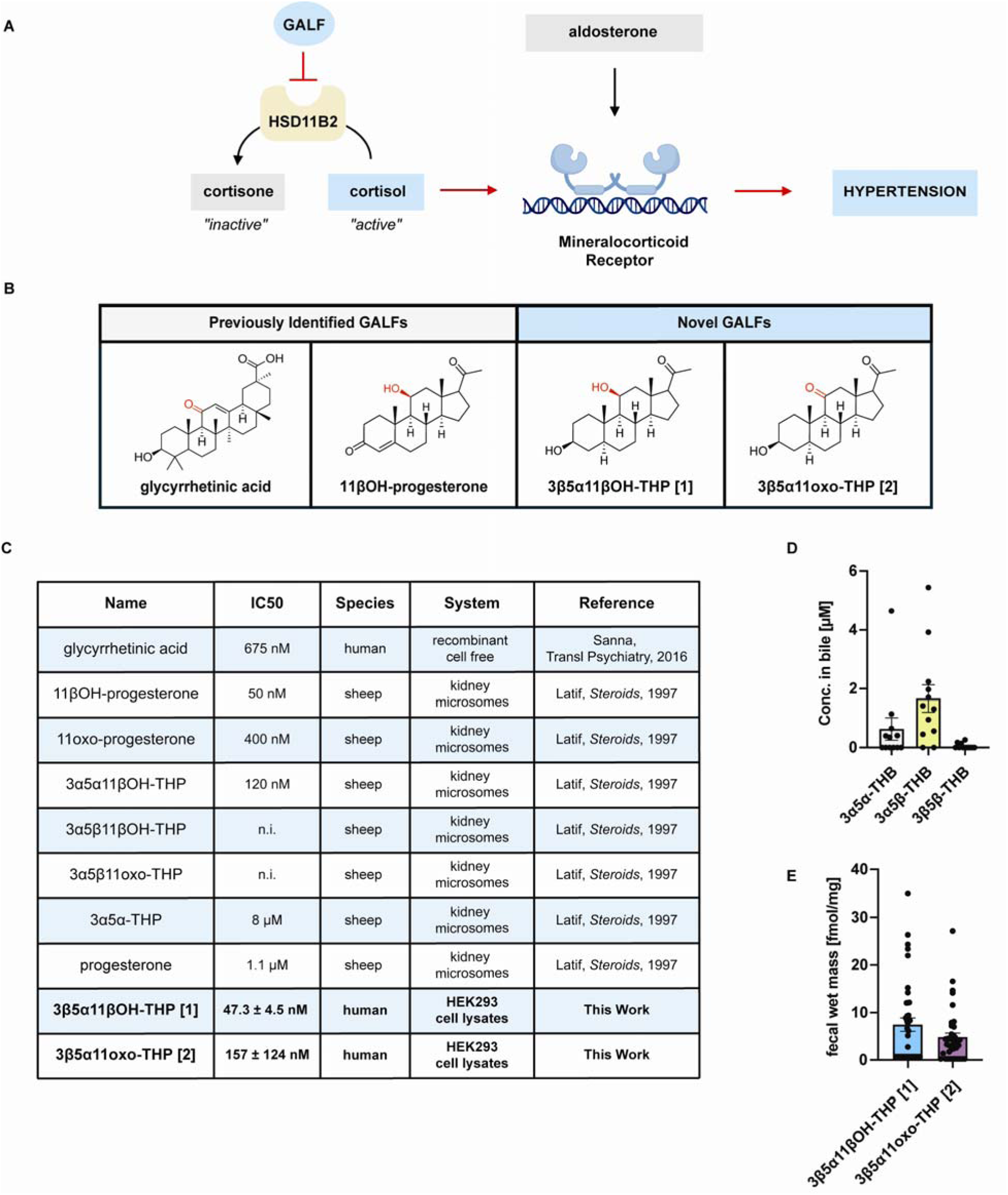
Identification of GALFs in human feces. (A) Inhibition of 11β hydroxysteroid dehydrogenase 2 (HSD11B2), a key enzyme in the mammalian cortisol shuttle, by glycyrrhetinic acid-like factors (GALFs), permits cortisol dependent activation of the mineralocorticoid receptor (MR) and promotes hypertension. (B) Structures of two previously identified GALFs, glycyrrhetinic acid and 11β hydroxyprogesterone, and GALFs identified in this work, 3β,11β dihydroxy 5α tetrahydroprogesterone (3β5α11βOH THP, **1**) and 3β hydroxy 11 oxo 5α tetrahydroprogesterone (3β5α11oxo THP, **2**). Red color highlights oxygenation at C11. (C) Reported HSD11B2 IC_50_ values for known GALF inhibitors and weak/non inhibitory steroids from prior studies as well as IC_50_ values for compounds **1** and **2** from this work. (D) Concentrations of upstream biliary corticoids tetrahydrocorticosterones (THBs) in human bile from patients with primary sclerosing cholangitis (PSC) and non-PSC controls, quantified by UHPLC-MS (n = 12 samples, one sample per patient). 3α5β-THB was the predominant metabolite in human bile. Bars show mean ± SEM. (E) Concentrations of **1** and **2** in healthy human fecal samples, quantified by UHPLC-MS (n = 39 samples). Values are reported as fmol/mg wet mass (approximately nM). Bars show mean ± SEM.

Gut bacteria can metabolize host-derived steroids in the gastrointestinal tract, generating structurally diverse metabolites. Steroid compounds such as tetrahydrocorticosterone (THB) and tetrahydrodeoxycorticosterone (THDOC) are present in bile at micromolar concentrations and enter the gastrointestinal tract via biliary excretion, where they can be further metabolized by gut bacteria.^47–49^ Gut bacteria are capable of diverse transformations of the steroid scaffold, including dehydroxylation, oxidation-reduction, and isomerization reactions that alter both functional groups and ring stereochemistry.^50–60^ Such modifications can profoundly influence receptor binding and enzyme inhibition. Consistent with this capacity, our prior work has shown that gut *Eggerthellaceae* convert host corticoids into progestin-like metabolites through 21-dehydroxylation.^50^ Altogether, these observations suggest that the microbiome may have the biochemical capability to generate steroid metabolites with GALF-like activity.

Here, we identify two steroid metabolites, 3β,11β-dihydroxy-5α-tetrahydroprogesterone (3β5α11βOH-THP, **1**) and 3β-hydroxy,11-oxo-5α-tetrahydroprogesterone (3β5α11oxo-THP, **2**), that potently inhibit human HSD11B2 (hHSD11B2), with nanomolar IC_50_ values measured in HEK-293 cell lysates. We further demonstrate inhibitory activity in more physiological systems, including intact SW-620 colon epithelial cells and patient-derived colon organoids that endogenously express hHSD11B2. Starting from the biliary corticoid 3α5β-tetrahydrocorticosterone (3α5β-THB), we show that gut bacteria can convert host corticoids into these HSD11B2 inhibitors through a series of transformations of the steroid scaffold, including 21-dehydroxylation, A-ring isomerization, and C11 oxidation. Both metabolites are detected in healthy human feces at concentrations sufficient to inhibit hHSD11B2 and depleted following antibiotic treatment, suggesting microbiome dependence. In addition, longitudinal analysis of human samples revealed that **1** is significantly elevated during pregnancy in stool, and both metabolites were higher in individuals with stage **2** hypertension-range blood pressure. Together, these findings identify microbiome-enabled steroid transformations that generate potent HSD11B2 inhibitors and suggest a potential link between gut microbial metabolism and MR activation that is relevant to human health.

## RESULTS

### Identification of GALF metabolites in human feces with conserved structural features

A defining structural feature of GALFs is 11-oxygenation of the steroid scaffold, which is required for inhibition of HSD11B2. For example, while 3α,11β-diOH-5α-THP (3α5α11βOH-THP), 11β-hydroxyprogesterone, and 11-oxoprogesterone have reported IC_50_ values of 120 nM, 50 nM, and 400 nM respectively, 3α-hydroxy-5α-pregnan-20-one (3α5α-THP) and progesterone are far less potent, with reported IC_50_ values of 8 µM and 1.1 µM respectively (**Fig. 1B & 1C)**.^40,61–63^ To determine whether potential GALF substrates are deposited into the gut, we analyzed enzymatically deconjugated human bile by UHPLC-MS for the presence of 11-oxygenated steroids and detected 3α5β-THB at low micromolar concentrations (**Fig. 1D**), consistent with prior reports.^47,64,65^ To identify potential bacterial products, we profiled healthy human fecal samples and identified several 11-oxygenated THP isomers, including the previously uncharacterized compounds 3β5α11βOH-THP (**1**) and 3β5α11oxo-THP (**2**), present at mean levels of 7 and 5 fmol/mg wet mass, respectively (fmol/mg wet mass is approximately equal to nM, **Fig. 1E** and **Supplementary Fig. 1A**). We confirmed the identity of these compounds using authentic standards, tandem MS, and spike-in experiments (**Supplementary Fig. 1B and 1C**).

Next, we evaluated the inhibitory activity of the identified metabolites in HEK-293 cell lysates stably overexpressing HSD11B2 (hHSD11B2). Both 3β5α11βOH-THP (**1**) and 3β5α11oxo-THP (**2**) potently inhibited enzyme activity, with IC_50_ values of 47.3 ± 4.5 nM and 157 ± 124 nM, respectively (**Fig. 2A**). In contrast, the structurally related 5β-reduced isomer 3α5β11βOH-THP exhibited substantially weaker inhibition, consistent with prior work (∼9.3 ± 1.1 μM).^61,62^ The known GALF 3α5α11βOH-THP was used as a positive control (IC_50_ 105.1 ± 14.8 nM; **Supplementary Fig. 1D**). Consistent with previous studies,^61–63,66^ these results indicate that 5α-reduced steroids, which adopt a planar A/B ring conformation, are more potent inhibitors of HSD11B2 than their 5β counterparts, which adopt a bent A/B ring conformation (**Fig. 2B**). These findings indicate that both 11-oxygenation and A-ring stereochemistry are key determinants of GALF activity.

**Figure 2.**
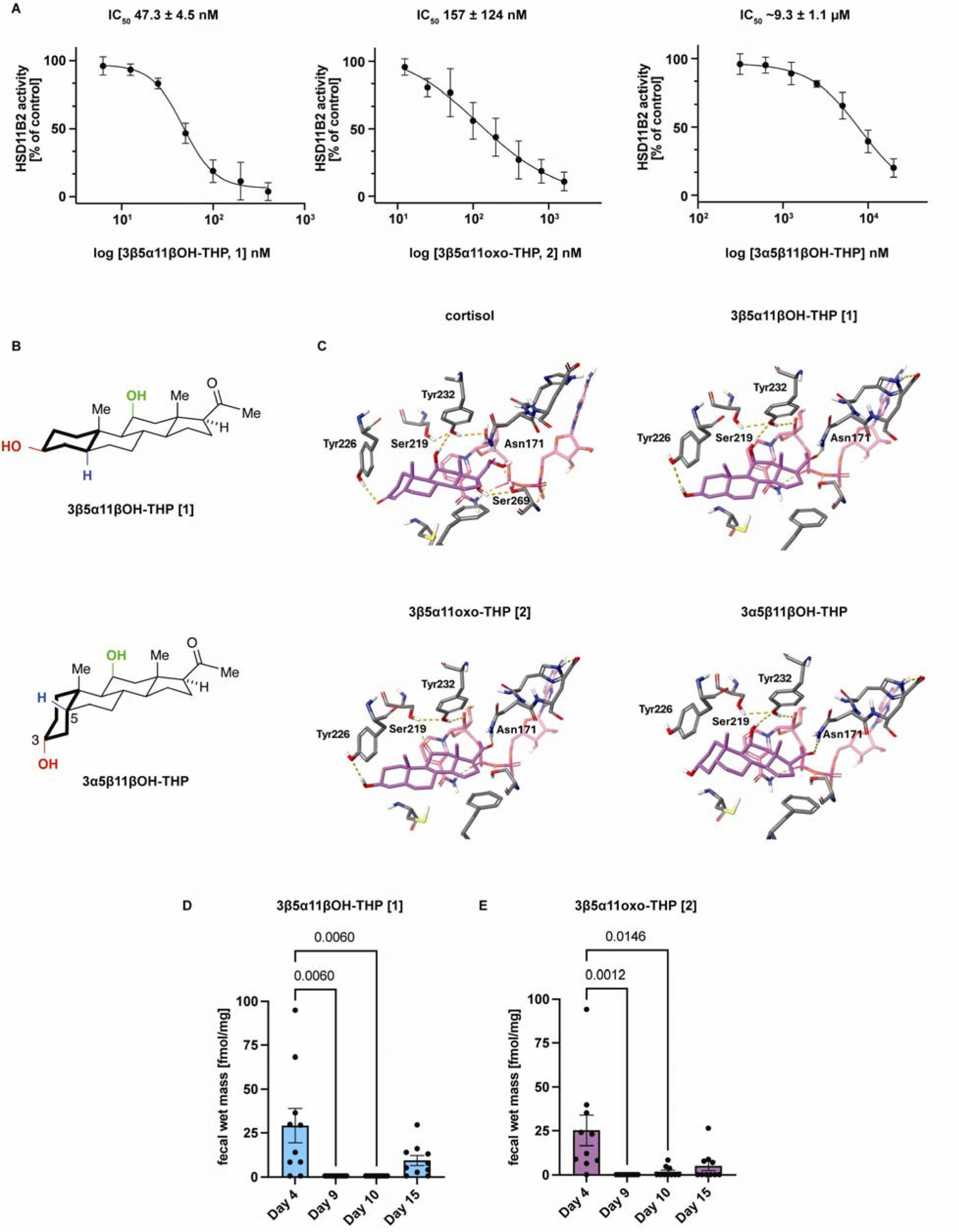
11-oxygenated 5α-reduced progestins are potent inhibitors of hHSD11B2 and are depleted in antibiotic-treated humans. (A) Concentration-response curves for inhibition of human HSD11B2 (hHSD11B2) in HEK-293 cell lysates by **1**, **2**, and 3α5β11βOH THP. Lysates were incubated with 40 nM cortisol, 10 nCi [³H]cortisol, 500 μM NAD⁺, and the indicated inhibitor concentrations for 10 min at 37°C. Conversion of cortisol to cortisone was normalized to 0.1% DMSO control. Data are mean ± SD from ≥3 biological replicates performed in technical duplicate; IC50 values were obtained by nonlinear regression. (B) 3D representations of **1** and 3α5β11βOH THP illustrating the planar 5α reduced (trans A/B ring) and bent 5β reduced (cis A/B ring) conformations, respectively. C3 oxygenation shown in red, 5 position stereochemistry shown in blue, and 11-oxygenation shown in green. (C) Cofolded models of hHSD11B2 showing docked poses of cortisol, **1**, **2**, and 3α5β11βOH THP. Dashed lines denote predicted hydrogen bonds between ligands and key active site residues (TYR226, TYR232, ASN171). (D-E) Longitudinal analysis of GALF metabolites in a controlled feeding antibiotic (Abx)/polyethylene glycol (PEG) microbiota purge study. Fecal levels of **1** (D) and **2** (E) were quantified by UHPLC–MS at the end of the baseline diet phase (day 4), immediately following the Abx/PEG phase (day 9), in the early recovery phase (day 10), and in the late recovery phase (day 15) (n=10 as in Tanes et al., 2021). Levels of both metabolites were significantly reduced by Abx/PEG treatment and began to rise by day 15. Statistical significance was assessed using a Friedman test followed by Dunn’s multiple-comparisons test. Bars show mean ± SEM.

To gain structural insight into GALF activity, we used cofolded modeling to compare the primary hHSD11B2 substrate cortisol with the identified GALFs. Cortisol, which contains a Δ4-5 double bond in the A-ring, formed hydrogen bonds with TYR226 through its C3-oxo group, with TYR232 through its C11 hydroxyl group, and with SER269 through its C17 hydroxyl group (**Fig. 2C**). The potent inhibitor 3β5α11βOH-THP [**1**] showed a similar fit within the hHSD11B2 binding pocket, forming hydrogen bonds with TYR226 through its C3 hydroxyl group, with TYR232 through its C11 hydroxyl group, and with ASN171 through its C20-oxo group. The less potent 11-oxo analogue 3β5α11oxo-THP [**2**] retained the interactions with TYR226 and ASN171, whereas its C11-oxo group formed a hydrogen bond with SER219 instead of TYR232. In contrast, the 5β-reduced isomer 3α5β11βOH-THP adopted a distinct binding pose, consistent with its non-planar cis A/B-ring conformation and did not show the C3 interaction observed for cortisol and the identified GALFs. However, it retained hydrogen bonds with TYR232 through its C11 hydroxyl group and with ASN171 through its C20-oxo group (**Fig. 2C**). Notably, the known inhibitor 3α5α11βOH-THP formed hydrogen bonds with TYR232 and ASN171 through its C11 hydroxyl and C20-oxo groups, respectively, while its C3 hydroxyl group interacted with CYS264 rather than TYR226 (**Supplementary Fig. 1E**). Together, these models identify conserved anchoring interactions at C11 and C20 and suggest that potency depends at least in part on how A-ring stereochemistry positions the C3 group: the more potent inhibitors engage either TYR226 or, for 3α5α11βOH-THP, CYS264, whereas the weak 5β isomer lacks an equivalent C3 interaction.

### Identified GALFs are reduced by microbiota depletion in human feces

To assess whether these metabolites are microbiome-dependent in vivo, we analyzed fecal samples from a published longitudinal controlled feeding study in which participants underwent microbiota depletion with antibiotics (Abx) and a polyethylene glycol (PEG) gut purge.^67^ Human subjects consumed a controlled omnivore diet, and treatment was divided into three phases: the dietary phase (days 1-5), the Abx/PEG gut microbiota purge designed to reduce bacterial load (days 6-8), and the recovery phase (days 9-15). Previous work with this cohort has confirmed bacterial load reduction during the Abx/PEG phase based on counting colony forming units (CFU), shotgun reads, and quantitative 16S rRNA gene copy number.^67,68^ GALF levels in feces were quantified during the dietary phase (day 4), immediately after microbiome depletion and in early recovery (days 9 and 10), and in late recovery (day 15). Levels of both metabolites **1** and **2** were significantly reduced following Abx/PEG treatment (day 4 vs day 9, **Fig. 2D and 2E**). During recovery, levels of **1** and **2** began to rise by day 15 (**Fig. 2D and 2E**). Additional C11 oxidized metabolites were also significantly depleted during the Abx/PEG phase (**Supplementary Fig. 2**). These findings indicate that depleting the microbiome substantially reduces the levels of GALFs **1** and **2** in the human gut and suggests that human gut bacteria may contribute to the production of these molecules in vivo.

### GALF metabolites exhibit murine vs human species-specific production and activity

We next examined whether GALF production and activity are conserved across humans and mice. In contrast to humans, where THB metabolites are present in bile at micromolar concentrations (**Fig. 1D**), THBs were detected at only nanomolar levels in mouse gallbladder samples (**Supplementary Fig. 3A**), representing a reduction in substrate availability of approximately three orders of magnitude. Consistent with this difference, both GALF metabolite levels were low to undetectable in mouse fecal samples, including in germ free (GF) mice colonized with human microbiota from pregnant and non-pregnant human donors (**Supplementary Fig. 3B and 3C**). Together, these data indicate that low availability of THB substrates in murine bile limits GALF levels in the mouse gut, even in the presence of a human microbiome.

**Figure 3.**
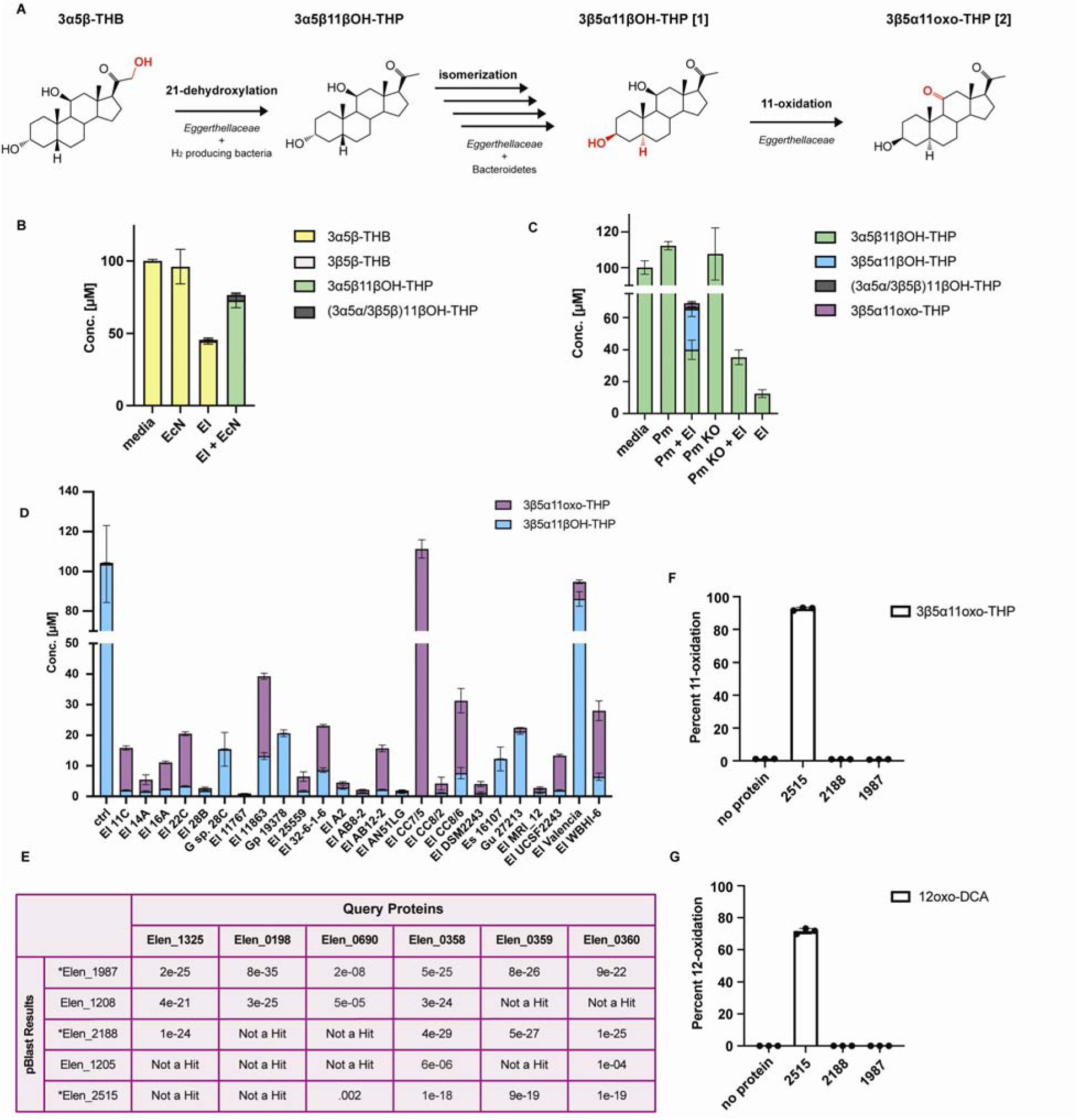
A multi-step interspecies bacterial enzymatic pathway converts host corticoids into GALF metabolites 1 and 2. (A) Proposed pathway for conversion of the biliary corticoid 3α5β-THB to GALF metabolites **1** and **2** by gut bacteria. Steps include 21-dehydroxylation of 3α5β-THB to yield 3α5β11βOH-THP, A-ring/C5 isomerization to generate **1**, and C11 oxidation to produce **2**. (B) Co-culture of *Eggerthella lenta* 14A (El) and *Escherichia coli* Nissle 1917 (EcN) converts 3α5β-THB to 3α5β11βOH-THP. Co-cultures, *E. lenta* 14A monocultures, and EcN monocultures were incubated with 100 μM 3α5β-THB for 48 h, and steroid products were quantified by UHPLC-MS (n = 3 biological replicates per condition). Only co-culture produced 3α5β11βOH-THP, consistent with 21-dehydroxylation activity requiring both partners. (C) Cooperative metabolism by *E. lenta* 14A and *Parabacteroides merdae* (Pm) converts 3α5β11βOH-THP to **1** and requires an A-ring isomerization gene cluster in *P. merdae*. Co-cultures of *E. lenta* 14A and wild-type *P. merdae* with substrate produced **1**, whereas *E. lenta* 14A or *P. merdae* monocultures, or co-culture with an isomerization-cluster knockout strain (*P. merdae* Δ04016-18, Pm KO), did not produce detectable **1** (n = 3 biological replicates per condition). (D) Screening of an *Eggerthellaceae* strain library revealed strain-level variation in C11 oxidation of **1**. Strains were incubated with 100 μM **1** for 72 h, and production of **2** was quantified by UHPLC-MS (n = 3 biological replicates per strain). Both *Eggerthella* and *Gordonibacter* species showed 11-oxidative activity with varying efficiencies. (El = *Eggerthella lenta*; Gp = *Gordonibacter pamelaeae*; Es = *Eggerthella sinensis*; G. sp = *Gordonibacter species*) (E) First step of the bioinformatic workflow used to identify candidate 11β-hydroxysteroid dehydrogenase (11βHSDH) genes in *E. lenta* DSM 2243. Previously characterized *E. lenta* hydroxysteroid dehydrogenases were used as queries in BLASTP searches against the DSM 2243 genome, yielding five gene candidates. Subsequent filtering for genes present across 11-oxidation-positive strains revealed three 11βHSDH gene candidates for *in vitro* testing: Elen_2188, Elen_2515, and Elen_1987 (see Supp Table 1). (F) The enzyme encoded by Elen_2515 exhibits 11βHSDH activity. Purified recombinant Elen_2188, Elen_2515, and Elen_1987 were incubated with **1** for 4 h at 37°C and conversion to **2** was quantified by UHPLC-MS (n = 3 independent reactions per enzyme, experiment was performed twice with similar results). Elen_2515 robustly oxidized **1** to **2**, whereas Elen_2188 and Elen_1987 showed no detectable activity. (G) Percent conversion of deoxycholic acid (DCA) to 12-oxoDCA by purified Elen_2515 under the same conditions as Fig. 3F, confirming its previously described 12α-hydroxysteroid dehydrogenase (12αHSDH) activity (n = 3 independent reactions per enzyme, experiment was performed twice with similar results).

We also observed species-specific differences in HSD11B2 inhibition. While 3β5α11βOH-THP (**1**) potently inhibited human HSD11B2 (hHSD11B2), it showed substantially weaker activity against mouse HSD11B2 (mHSD11B2) (IC_50_ = 1.32 ± 0.19 µM**; Supplementary Fig. 3D**). Cofolded modelling revealed differences in ligand accommodation between hHSD11B2 and mHSD11B2. While the natural murine substrate corticosterone was well accommodated within the mHSD11B2 binding pocket, **1** could not be docked. The compound was therefore positioned manually based on the corticosterone orientation. In this orientation, the C3 hydroxyl group showed a steric clash with MET347, the C11 hydroxyl group with TYR232, and the C20-oxo group with ASN171 (**Supplementary Fig. 3E**). These clashes indicate less favorable accommodation of **1** within the murine binding pocket. However, because the compound was positioned manually, its actual binding orientation and potential steric clashes remain uncertain. These findings indicate species-specific differences in binding-pocket geometry that may impair inhibitor binding to mHSD11B2.

### Gut bacteria produce GALFs through a multi-step enzymatic pathway

Gut bacteria are known to metabolize host-derived corticoids. Prior work from our lab demonstrated that *Eggerthellaceae*, together with hydrogen-producing partner species, convert tetrahydrodeoxycorticosterone (THDOC) to tetrahydroprogesterone (THP) via 21-dehydroxylation.^50^ In addition, 3α5β-THB, an abundant 11-oxygenated corticoid present in human bile (**Fig. 1D**), represents a potential substrate for downstream GALF production. We therefore hypothesized that gut bacteria convert 3α5β-THB to GALFs through a series of transformations: 21-dehydroxylation to generate 3α5β11βOH-THP, A-ring isomerization at C3 and C5 to produce 3β5α11βOH-THP (**1**), and oxidation at C11 to yield 3β5α11oxo-THP (**2**) (**Fig. 3A**).

To test the first step, we co-cultured the 21-dehydroxlyating strain *Eggerthella lenta* 14A with the hydrogen-producer *E. coli* Nissle in the presence of 3α5β-THB and observed conversion to 3α5β11βOH-THP, demonstrating 21-dehydroxylation activity. Neither *E. lenta* 14A alone nor *E. coli* Nissle alone produced 3α5β11βOH-THP, consistent with our prior work showing that neither bacterium can perform 21-dehydroxylation on its own (**Fig. 3B**).^50^

We next examined A-ring isomerization. Previous work from our lab and others identified a biosynthetic gene cluster in gut Bacteroidota species encoding enzymes that mediate sequential C5 and C3 transformations, including a 5β-reductase, a 5α-reductase, and a 3β-hydroxysteroid dehydrogenase (3βHSDH).^60,69^ However, this pathway has been characterized only for bile acid substrates, and Bacteroidota lack a 3αHSDH required for the initial oxidation step of the steroidal core. We therefore combined *E. lenta* 14A, which encodes 3αHSDH activity,^70,71^ with *Parabacteroides merdae*, a representative Bacteroidota strain harboring this gene cluster. Co-culture of these species resulted in conversion of 3α5β11βOH-THP to 3β5α11βOH-THP (**1**). (**Fig. 3C**). Monoculture with either *E. lenta* 14A or *P. merdae* with substrate resulted in no detectable isomerized product. *E. lenta* 14A monoculture with 3α5β11βOH-THP resulted in some conversion to an oxidized product with an m/z consistent with the intermediate 3oxo-5β11βOH-THP, suggesting that this bacterium performs the initial 3α-oxidation step (**Supplementary Fig. S4**). Co-culture of *E. lenta* 14A with *P. merdae* Δ04016-18 (Pm KO), a genetic deletion strain that lacks the isomerization gene cluster,^60^ resulted in no formation of isomerized product, indicating that this cluster is required for isomerization by *P. merdae* (**Fig. 3C**). Overall, these results demonstrate that A-ring isomerization of corticoid-derived substrates can be achieved through cooperative interspecies bacterial metabolism.

Next, we investigated formation of the second GALF metabolite 3β5α11oxo-THP (**2**) via C11 oxidation. Low levels of **2** were observed in *E. lenta* 14A and *P. merdae* co-cultures, suggesting that this activity is encoded by these community members. Given prior evidence that *Eggerthellaceae* encode HSDHs that act on steroid substrates,^50,70–72^ we screened an *Eggerthellaceae* strain library for 11-oxidative activity using **1** as the substrate. Multiple strains converted **1** to **2**, indicating that this activity is broadly distributed within *Eggerthella* and *Gordonibacter* species (**Fig. 3D**).

To identify candidate enzymes responsible for C11 oxidation, we performed a sequence-guided search of the *E. lenta* DSM 2243 type strain genome using six previously characterized *E. lenta* HSDHs as query sequences. This analysis yielded a set of putative HSDH-encoding genes (**Fig. 3E**). We then applied a second filtering step, retaining only candidates present in all strains exhibiting 11-oxidative activity (**Supplementary Table 1**). This approach identified three candidate genes, Elen_1987, Elen_2515, and Elen_2188. Elen_2515 has been previously characterized as a 12α-hydroxysteroid dehydrogenase (12αHSDH) that catalyzes C12 oxidation of bile acids such as deoxycholic acid (DCA).^70,72^ Given the close proximity of the C11 and C12 positions on the steroid C-ring, we hypothesized that Elen_2515 may also exhibit activity at the C11 position. Recombinant expression and biochemical characterization of these three candidates revealed that Elen_2515 catalyzes oxidation of **1** to **2**, as detected by UHPLC-MS, while Elen_2188 and Elen_1987 displayed no activity (**Fig. 3F**). Consistent with prior reports, Elen_2515 also oxidized deoxycholic acid to 12-oxoDCA (**Fig. 3G**).^70,72^ Notably, we find that Elen_2515 catalyzes C11 oxidation, identifying it as a bacterial 11β-hydroxysteroid dehydrogenase (11βHSDH) with dual 11β- and 12α-oxidative activity. Overall, these results reveal a multi-step pathway for GALF production from host-derived corticoids by human gut bacteria.

### Bacterially produced GALFs inhibit endogenous hHSD11B2 in human colonic cells

To assess whether these metabolites inhibit endogenous HSD11B2 activity in a cellular context, we measured the conversion of cortisol to cortisone in intact SW-620 colon epithelial cells, which endogenously express hHSD11B2. Treatment with 3β5α11βOH-THP (**1**) reduced cortisol-to-cortisone conversion in a concentration-dependent manner, with ∼40% inhibition at 20 nM and near-complete inhibition at 1 µM (**Fig. 4A**). In contrast, 3α5β11βOH-THP showed weak inhibition as expected, producing ∼30% inhibition at 1 µM (**Fig. 4A**). Consistent with these results, **1** exhibited an IC_50_ of 30.2 ± 1.1 nM in this intact cell system (**Fig. 4B**). In line with the lysate-based assays, 3β5α11oxo-THP (**2**) demonstrated an IC_50_ of 54.3 ± 11.1 nM (**Fig. 4C**). These values provided a benchmark for evaluating whether GALF concentrations measured in human feces reached the range capable of inhibiting HSD11B2 in intact colonic cells. Notably, neither metabolite exhibited detectable cytotoxicity under any of these conditions (**Supplementary Fig. 5A and 5B**).

**Figure 4.**
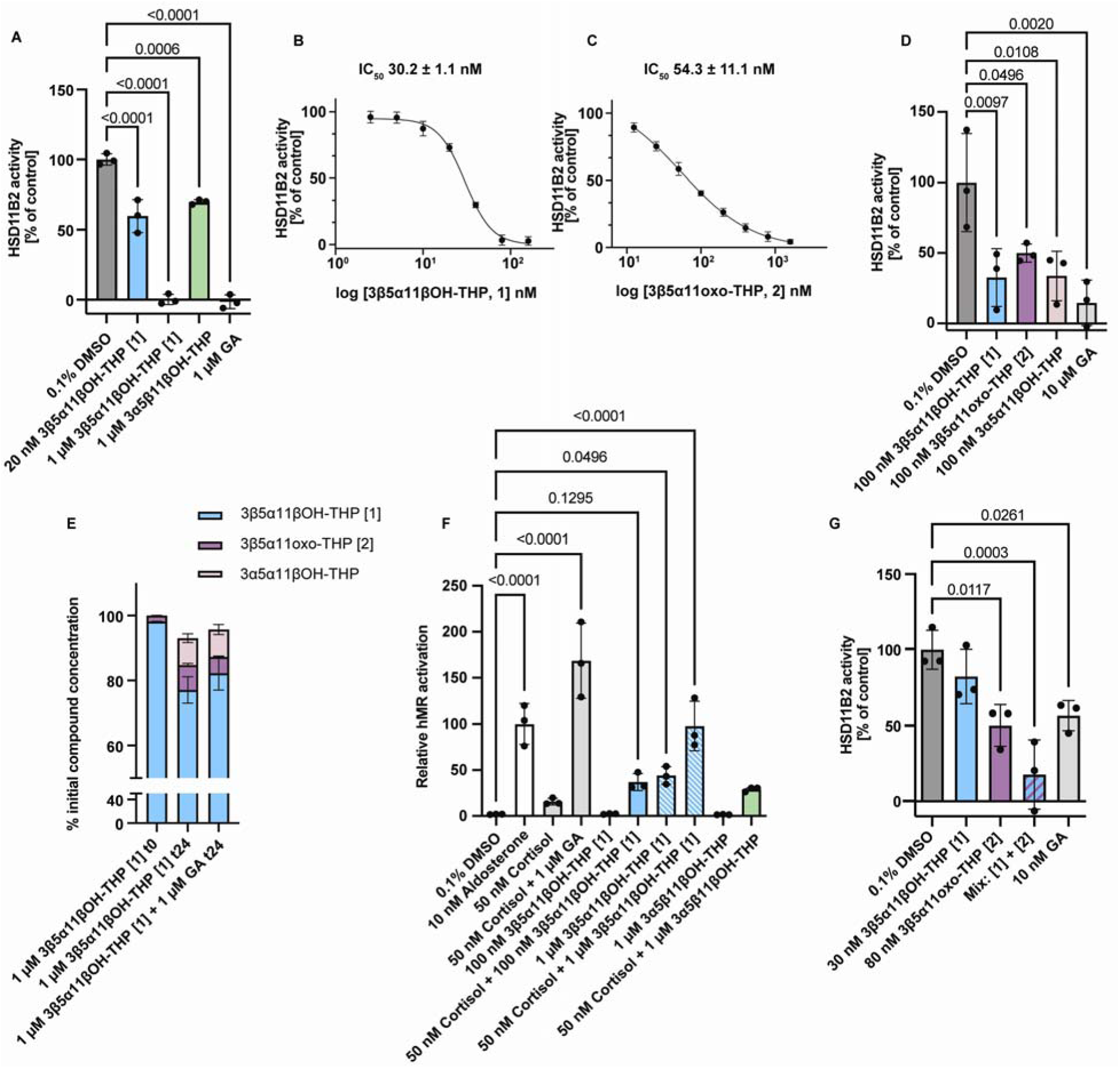
GALF metabolites 1 and 2 inhibit endogenous hHSD11B2 resulting in mineralocorticoid receptor activation in human colonic cells and organoids. (A) Compound **1** inhibits endogenous hHSD11B2 in intact SW 620 colon epithelial cells. Cells were treated with 40 nM cortisol and 10 nCi [³H]cortisol in the presence of the indicated concentrations of **1**, 3α5β11βOH THP, or 1 μM glycyrrhetinic acid (GA; positive control) for 4 h at 37°C. HSD11B2 activity was calculated as cortisone/(cortisol + cortisone) and normalized to 0.1% DMSO control (n = 3 independent experiments, each in technical duplicate). Bars show mean ± SD. (B-C) Concentration-response curves and IC₅₀ values for **1** (B) and **2** (C) in intact SW 620 cells. Data are mean ± SD from ≥3 independent experiments performed in technical duplicate; curves were fit by nonlinear regression. (D) Compounds **1** and **2** inhibit HSD11B2 activity in intact colon organoids. Organoids derived from donor P1 were treated with 40 nM cortisol, 10 nCi [³H]cortisol, and 100 nM compound **1**, 100 nM compound **2**, 100 nM 3α5α11βOH THP, or 10 μM GA for 6 h at 37°C. HSD11B2 activity was quantified as in (A) (n = 3 independent experiments, each in technical duplicate). (E) Compound **1** is not readily oxidized in SW 620 cells. Cells were incubated with **1** alone (1 µM) for 0 h or 24 h or **1** (1 µM) and 1 µM GA for 24 h; supernatants were analyzed by UHPLC-MS/MS to quantify formation of **2**. Data are expressed as percent of initial compound (n = 3 biological replicates per time point). (F) Compound **1** activates hMR in the presence of cortisol. MR transactivation in V79 cells transiently overexpressing human MR (hMR), HSD11B2, an MMTV LacZ reporter, and a CMV driven luciferase transfection control (cluc). Aldosterone (10 nM) and cortisol (50 nM) plus GA (1µM) were used as positive controls (n = 3 independent experiments, each in technical duplicate). (G) GALFs **1** and **2** cooperatively inhibited hHSD11B2 in HEK293 cells stably overexpressing hHSD11B2. Cells were incubated with 40 nM cortisol, 10 nCi [³H]cortisol, and 30 nM **1**, 80 nM **2**, a combination of **1** and **2**, or 10 nM GA for 10 min. HSD11B2 activity was calculated as in Fig. 4A and normalized to 0.1% DMSO control. Combined treatment with **1** and **2** resulted in additive inhibitory effects (n = 3 independent experiments in technical duplicate). All data are presented as mean ± SD. Statistical analyses were performed by one way ANOVA followed by Dunnett’s multiple comparison test unless otherwise indicated. No cytotoxicity was detected under any condition (see Figure S5 and S6).

**Figure 5:**
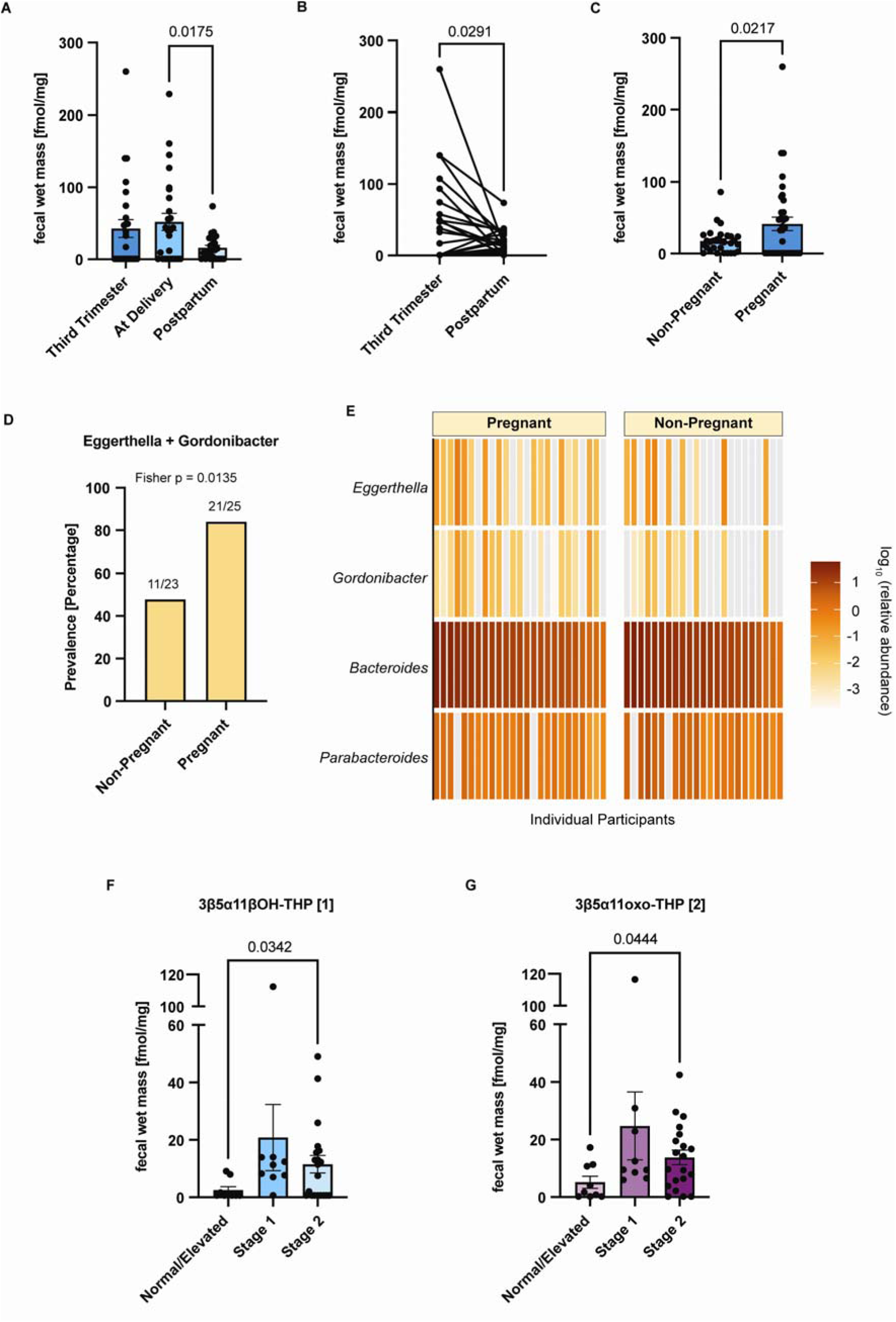
GALF metabolites are increased during pregnancy and in individuals with stage 2 hypertension-range blood pressure. (A) Longitudinal fecal levels of **1** in a Finnish pregnancy cohort. Fecal samples collected in the third trimester, at delivery, and 11-18 months postpartum were analyzed by UHPLC-MS (n = 26 samples per time point, bars show mean ± SEM; fmol/mg wet mass is approximately equal to nM). One-way repeated measures ANOVA with Geisser-Greenhouse correction and Tukey’s multiple comparisons. (B) Paired analysis of fecal **1** levels in individuals sampled during late pregnancy and postpartum (n = 26 matched pairs). Lines connect samples from the same individual. (C) Comparison of fecal **1** levels between pregnant individuals in the Finnish cohort (third trimester from samples with longitudinal data (n=26) with additional third trimester samples without longitudinal data (n=9); total (n = 35)) and non pregnant Finnish controls (n = 30). (D) Prevalence of *Eggerthella* and *Gordonibacter* genera in non-pregnant and pregnant stool samples. eleven out of 23 non-pregnant samples contained these genera, and 21 out of 25 pregnant samples contained these genera. Two-sided fisher exact test was performed (p= 0.0135). (E) Select metagenomic abundance of *Eggerthella*, *Gordonibacter*, *Bacteroides*, and *Parabacteroides* genera in stool samples from pregnant (n= 25) and non-pregnant (n= 23) Finnish donors. One pregnant donor not graphed due to lack of metagenomic information. (F and G) Concentrations of **1** and **2** are increased in patients with stage 2-range blood pressure compared to normal/elevated blood pressure in a Boston-based cohort of patients with colorectal adenomas (n = 38). Normal/elevated blood pressure (systolic <130 mmHg & diastolic <80 mmHg), stage 1 hypertension-range blood pressure (systolic 130-139 mmHg or diastolic 80-89 mmHg), stage 2 hypertension-range blood pressure (systolic ≥140 mmHg or diastolic ≥90 mmHg). Statistical significance was assessed using Brown-Forsythe and Welch ANOVA followed by Dunnett’s T3 multiple-comparisons test. All metabolites were quantified by UHPLC-MS and are reported as fmol/mg wet mass. All data are presented as mean ± SEM. Paired t tests were used for paired longitudinal analyses; unpaired t tests were used for unpaired analyses.

We next evaluated inhibition in a more physiologically relevant system using patient-derived human colon organoids. HSD11B2 expression was confirmed by RT-qPCR and Western blot (**Supplementary Fig. 5C and 5D**). Consistent with results in SW-620 cells, **1** potently inhibited cortisol-to-cortisone conversion, reducing activity to ∼30% at 100 nM (**Fig. 4D**). Compound **2** also inhibited HSD11B2, though less effectively. The positive control, 3α5α11βOH-THP, also produced a similar inhibition profile. No cytotoxicity was observed (**Supplementary Fig. 5E**).

The 11β-hydroxy group of corticosterone and cortisol can be oxidized to a carbonyl by HSD11B2. Therefore, we next asked whether **1** is not only an inhibitor but also a substrate of this enzyme. UHPLC-MS/MS analysis of SW-620 cells treated with **1** revealed limited conversion to the corresponding 11-keto product **2**, with only ∼6% net formation after 24 h (**Fig. 4E**). This conversion was suppressed by GA supplementation. In contrast, ∼70% conversion of cortisol to cortisone occurred under the same conditions, and this transformation was blocked by GA addition (**Supplementary Fig. 5F**). No toxicity was observed under any of these conditions (**Supplementary Fig. 5G**). These data indicate that while the 11β-hydroxy GALF **1** can serve as a substrate for HSD11B2, turnover is inefficient relative to cortisol. Together, these results demonstrate that bacterially derived GALFs potently inhibit endogenous HSD11B2 activity in human colonic systems while **1** acts as a poor substrate for this enzyme.

### GALF metabolites cause MR activation in the presence of cortisol

We next tested whether GALF metabolites disrupt the gatekeeper function of HSD11B2, which normally prevents cortisol from activating the mineralocorticoid receptor (MR) by converting it to inactive cortisone.^33,38,40,44^ In V79 cells transiently over-expressing hMR and HSD11B2, 10 nM aldosterone robustly activated MR, whereas 50 nM cortisol failed to do so due to HSD11B2-mediated inactivation (**Fig. 4F**). Co-treatment with cortisol and the known HSD11B2 inhibitor GA restored MR activation, validating the assay.

Strikingly, co-treatment with cortisol (50 nM) and **1** (100 nM) also restored MR activation similarly to GA. This effect was concentration-dependent, with 1 μM of **1** producing a stronger response (**Fig. 4F**). In contrast, at 1 µM, 3α5β11βOH-THP produced only mild inhibition of HSD11B2 in the presence of cortisol (50 nM) resulting in modest MR activation, as expected. Consistent with this mechanism, **1** alone did not activate MR at 100 nM, whereas modest activation was observed at 1 μM, suggesting a secondary, potentially direct effect on hMR at higher concentrations. In the absence of HSD11B2, **1** directly activated hMR in a transactivation assay, although with substantially lower potency than cortisol (**Supplementary Fig. 6A and 6B**). These effects were modest and only apparent in the absence of HSD11B2, conditions that are unlikely to be physiologically relevant in colonic cells, indicating that direct MR activation likely represents a secondary mechanism relative to HSD11B2 inhibition. We confirmed that the treatments used in these assays were not cytotoxic (**Supplementary Fig. 6C and 6D**).

Finally, we assessed whether multiple microbiome-derived GALFs can act cooperatively to inhibit HSD11B2. In HEK-293 cells stably overexpressing hHSD11B2, low concentrations of **1** (30 nM) and **2** (80 nM) produced partial inhibition individually, whereas co-treatment resulted in substantially greater inhibition (∼75–80% reduction in activity) (**Fig. 4G**). GA was included as a reference control at 10 nM, a concentration close to its reported IC₅₀ of 12 ± 3 nM.^73^ Consistent with previous results, GA reduced HSD11B2 activity by approximately 50%, confirming assay performance. Together, these results demonstrate that bacterially produced GALFs block HSD11B2-mediated protection and enable cortisol-dependent MR activation, linking microbial metabolism to host steroid receptor signaling in the colon.

### 3β5α11βOH-THP is elevated during pregnancy in humans

Because both host steroid availability and gut microbial composition change during pregnancy, we next asked whether GALF levels also vary across pregnancy and postpartum periods. To address this question, we analyzed longitudinal stool samples from a Finnish pregnancy cohort using UHPLC-MS. This dataset included 117 stool samples collected across pregnancy (third trimester), delivery, and 11-18 months postpartum, including a subset of 26 longitudinally sampled individuals, along with 30 female non-pregnant controls (**Data Table 5**).^74,75^ Across this cohort, we detected a range of C11 steroid metabolites. Consistent with bile metabolite profiles, 3α5β-THB emerged as a dominant upstream metabolite, while multiple downstream 11β-hydroxy and 11-oxo derivatives were observed in stool (**Supplementary Fig. 7A)**. Expanding on prior findings in blood,^76–78^ we also detected progesterone and the estrogen metabolites estradiol and estrone in stool,^64,78^ with levels declining sharply postpartum (**Supplementary Fig. 7B**).

Within this metabolic network, both GALF metabolites were readily detected in human feces (**1**, mean during pregnancy: 43 fmol/mg, or approximately 43 nM; **2**, mean during pregnancy: 18 fmol/mg, or approximately 18 nM, **Fig. 5A** and **Supplementary Fig. 8A**). These concentrations overlap the 30.2 nM IC_50_ measured in intact human colonic cells, and levels in many individuals exceeded this value (**Fig. 4B** and **5A**), indicating that pregnancy-associated fecal concentrations of **1** reach a range capable of inhibiting HSD11B2 in cells. Levels of **1** were significantly elevated at delivery compared to postpartum timepoints (**Fig. 5A**), and paired analyses confirmed a decline after pregnancy (**Fig. 5B**). In contrast, **2** was consistently detected but did not vary significantly across timepoints (**Supplementary Fig. 8A**). Comparison to an independent non-pregnant Finnish cohort further confirmed that **1** was enriched during the third trimester (**1**, mean in nonpregnant cohort: 18 ± 3 SEM fmol/mg, **Fig. 5C**), whereas **2** was only marginally higher during pregnancy and did not reach statistical significance (**Supp Fig. 8B**). Because *Eggerthella* and *Gordonibacter* can perform steps in GALF biosynthesis, we examined their representation in this cohort. At least one of these genera was detected in a greater proportion of pregnant than non-pregnant samples (84% versus 48%; **Fig. 5D**), and both genera were numerically more abundant in pregnant samples (**Fig. 5E**), consistent with our prior work.^50^ Together, these results identify **1** as the GALF most strongly associated with pregnancy and show that pregnancy is also accompanied by enrichment of bacterial genera capable of participating in GALF biosynthesis.

### Fecal GALF levels are associated with elevated blood pressure in humans

Hypertension can arise from identifiable medical conditions, but primary hypertension has no single known cause and likely reflects both genetic and environmental influences, including diet and potentially the gut microbiome.^7,40,41,45,46,61^ Because the relationship between the GALFs identified here and human blood pressure had not previously been examined, we quantified **1** and **2** in 53 stool samples from patients with colorectal adenomas whose blood pressure was measured during the same baseline study visit, shortly before stool collection. We excluded 15 patients receiving antihypertensive medications and classified the remaining 38 based on this single blood pressure measurement as having normal/elevated, stage 1 hypertension-range, or stage 2 hypertension-range blood pressure. Both GALFs showed higher mean levels in patients with stage 1 or stage 2 hypertension-range blood pressure than in the normal/elevated group. These differences reached statistical significance in the stage 2 group (**Fig. 5F** and **5G**). The concentrations observed in these groups overlapped the nanomolar range that inhibited HSD11B2 in intact human colonic cells, supporting their potential physiological relevance. Because classification was based on a single blood pressure measurement rather than a clinical diagnosis of hypertension, these results require confirmation in larger cohorts with repeated measurements. Nonetheless, these findings provide initial evidence that fecal GALF levels are associated with elevated blood pressure in humans.

## DISCUSSION

In this study, we demonstrate that gut bacteria produce GALFs that can modulate host corticosteroid signaling. We also show that gut bacteria convert host-derived corticoids into GALFs through a multi-step enzymatic pathway whose genes are encoded in bacteria from different phyla. These metabolites potently inhibit human HSD11B2, blocking its gatekeeper function and enabling cortisol-dependent activation of MR. Notably, GALFs can act cooperatively, suggesting that ensembles of structurally related microbial metabolites can achieve physiologically relevant inhibition even at submaximal individual concentrations. These data support a model in which structurally related GALFs act in concert to suppress HSD11B2 activity, consistent with a “cloud” mechanism of inhibition.^40^ In humans, levels of these metabolites are reduced following antibiotic treatment, supporting a microbial contribution to their production. We also demonstrate that GALF **1** was elevated in feces during pregnancy, a physiological state in which corticosteroid signaling is tightly regulated. Finally, GALF levels were higher in individuals with stage 2 hypertension-range blood pressure, providing initial evidence of an association between these metabolites and elevated blood pressure in humans.

Our data show that GALF activity depends on both a planar A/B-ring configuration and 11-oxygenation, a structural orientation that facilitates productive engagement with the HSD11B2 active site. This finding is consistent with prior work showing that potent inhibitors of 11β-dehydrogenases are enriched for 11- and 7-oxygenated steroid scaffolds, including Δ4-3-oxo steroids such as cortisol and corticosterone, the primary substrates in humans and rodents, respectively.^40,61–63^ Together, these results extend established structure-activity relationships to bacterially produced metabolites.

The colon represents a key interface for MR-mediated epithelial sodium transport, potassium excretion and blood pressure regulation.^16,18,21,79,80^ Within this context, our data in colonic epithelial cells and patient-derived organoids demonstrate that GALFs inhibit HSD11B2 and enable cortisol-dependent MR activation, supporting a model in which locally produced microbial metabolites modulate host corticosteroid signaling. Importantly, fecal concentrations of **1** in humans overlapped the nanomolar concentration-response range measured in intact human colonic cells, and concentrations in some subjects exceeded the corresponding IC_50_ values. Although fecal measurements do not establish concentrations at the epithelial surface, these concentrations support the physiological plausibility that locally produced GALFs inhibit colonic HSD11B2 and enhance glucocorticoid-dependent MR signaling in at least a subset of individuals.

In addition to elucidating effects on host cells, we identify a bacterial enzyme with 11β-hydroxysteroid dehydrogenase activity. While bacterial steroid metabolism has been reported, 11-oxidative activity has not previously been described.^55–59^ The observation that **1** is a poor substrate for hHSD11B2, together with robust 11βHSDH activity in *Eggerthellaceae*, supports a microbial origin for 11-oxo THP metabolites. Notably, while Elen_2515 has previously been characterized as a 12αHSDH,^70,72^ our findings reveal that it also catalyzes C11 oxidation. This dual activity is consistent with prior examples of hydroxysteroid dehydrogenases that act at multiple positions, including mammalian HSD11B1, which reduces both 11-oxo glucocorticoids and 7-oxo bile acids,^81–85^ and a bacterial enzyme from *Comamonas testosteroni*, which exhibits 3β/17β-HSDH activity and catalyzes reactions at both the C3 and C17 positions of the steroid core.^86–88^ Together, these findings reveal that some HSDH enzymes are catalytically flexible and expand the known repertoire of microbiome-encoded steroid transformations.

Substantial human versus mouse species differences in GALF activity were observed, underscoring limitations of mouse models for studying these metabolites. Consistent with prior reports of interspecies variation in HSD11B2 inhibition by GA metabolites, oxysterols, and azole compounds,^73,89,90^ we found that bacterial GALFs exhibited reduced inhibitory effects on mHSD11B2. In parallel, GALF metabolite levels were low to undetectable in mice, even following human FMT. These findings suggest that both enzyme structure and metabolic differences contribute to species specificity and highlight the challenges of modeling microbiome-dependent steroid metabolism in rodents.

Our results further suggest that GALFs may be relevant to human blood pressure regulation, including during pregnancy. Dysregulation of the HSD11B2–MR axis is linked to hypertension and hypokalemia and may contribute to pregnancy-associated disorders such as preeclampsia, which is characterized by high blood pressure, proteinuria, and renal insufficiency.^3,91–97^ Fecal levels of **1** were elevated during pregnancy and declined postpartum. In a separate human cohort, both fecal GALFs were higher in individuals with stage 1 or stage 2 hypertension-range blood pressure than in the normal/elevated reference group, with statistically significant differences in the stage 2 group. The overlap between these human fecal concentrations and concentrations that inhibited HSD11B2 in intact colonic cells strengthens the biological plausibility of these associations. Although prior studies have shown that corticosteroid levels increase during pregnancy,^96^ our data suggest that changes in bacterially produced metabolites may also contribute to altered corticosteroid signaling. Increased availability of host steroid substrates may contribute to the pregnancy-associated increase in GALFs. Historical analyses reported higher mean concentrations of both 3α5α-THB and 3α5β-THB in bile collected during late pregnancy than in non-pregnant women.^47,65^ Together with our finding that *Eggerthella* and *Gordonibacter* are more prevalent during pregnancy, these observations suggest that changes in both host substrate availability and microbial composition could favor GALF production. Determining the relative contributions of these factors will require longitudinal studies combining GALF and host steroid measurements with bacterial gene abundance and expression across pregnancy. While a causal role for GALFs in preeclampsia or hypertension remains to be established, our findings provide a foundation for future studies examining microbiome-dependent steroid metabolism in blood pressure regulation and maternal health.

### Limitations of the study

First, while we define a biosynthetic pathway for GALF production, it is possible that additional microbial taxa and microbial and host enzymes contribute to this metabolism in vivo. Future studies integrating metagenomic and metatranscriptomic analyses with GALF and host steroid profiling will be required to determine how changes in microbial pathway abundance and substrate availability contribute to GALF production during hypertension-associated conditions. Second, although our cellular and organoid models support a role for GALFs in inhibiting HSD11B2 and modulating MR signaling, testing in additional human-relevant systems will be important to establish in vivo effects. This task is currently made more difficult by our finding of species-specific differences. Because both metabolite production and enzyme sensitivity are substantially lower in mice than humans, standard mouse models are of limited utility, highlighting the need for alternative approaches to study the biological effects of microbially produced steroids in humans.

Despite these limitations, these results establish a mechanistic link between bacterial steroid metabolism and host endocrine signaling. Furthermore, they suggest that microbial modulation of the HSD11B2-MR axis may contribute to human physiology and disease, particularly in the context of pregnancy.

## ACKNOWLEDGMENTS

We thank the Alm lab at MIT for providing feces from non-pregnant male and female donors and Joshua Korzenik for providing human bile samples. We thank Melissa Tran in the Devlin and Huh lab at HMS for providing gallbladders from mouse samples. We thank the Turnbaugh lab for providing the Actinobacteria strain library. We thank the Swiss National Science Foundation for supporting this project (No 310030_214978 to AO). We thank Dr. Martin Smieško for help with computational modeling. We also thank Prof. Dr. Salvatore Piscuoglio, University Hospital Basel for providing colon organoids and for access to protocols and training. Y.B. was supported by a grant of the Boston-Korea Innovative Research Project through the Korea Health Industry Development Institute (KHIDI), funded by the Ministry of Health & Welfare, Republic of Korea (RS-2024-00403047), and the Strategic Hub for International Research Collaboration project of Seoul National University. This work was supported by National Institutes of Health (NIH) grants R35 GM128618 (A.S.D.), R01 CA243454 (A.T.C.), and R35 CA253185 (M.S.), the Harvard Medical School - Seoul National University Hospital - Seoul National University College of Medicine Collaborative Research Program grant (A.S.D.), and the Armenise Harvard Foundation (A.S.D.). The Finnish EDIA study was supported by the National Institutes of Health (P30 DK043351), the Academy of Finland Centre of Excellence in Molecular Systems Immunology and Physiology Research 2012–2017 (250114; M.K.), and the Medical Research Funds of Tampere and Helsinki University Hospitals (M.K.).

## AUTHOR CONTRIBUTIONS

S.O.M., J.T.W., P.S.P. and M.L. conducted the experiments and analyzed the data. A.S.D., A.O., J.T.W., and S.O.M. conceived the project and A.S.D., A.O., J.T.W., S.O.M., P.S.P., and M.L. designed the experiments. J.T.W. performed the bacterial culture, bacterial cloning and protein expression experiments, developed UHPLC mass spectrometry methods to detect steroid metabolites in bacterial and biological samples, performed mass spectrometry experiments and analysis, and tested culture conditions. S.O.M. performed cell and organoid culturing, cell lysate and intact cell experiments, organoid experiments, performed and analyzed UHPLC-MS/MS experiments, and tested cell culture conditions. P.S.P. contributed to cell culture experiments and conducted additive effects experiment. M.L. performed metagenomic data analysis under the supervision of V.T. J.P. contributed to mass spectrometry experiments and analysis of biological samples under the supervision of A.S.D and Y.B. I.Z. provided mouse samples and conducted FMT mouse experiment under the supervision of J.R.H., and M.D.M. contributed to FMT mouse experiment. M.G.M. provided colorectal adenoma stool sample metadata, and T.H. provided pregnant and nonpregnant stool sample metadata. J.E.B. aided in metagenomic analysis. D.V.W developed the UHPLC-MS/MS method and extraction procedures to detect steroids in cell culture experiments. R.E.V.D performed cofolded docking experiments. M.K. provided pregnant and nonpregnant stool samples. M.S, D.A.D, A.T.C, and P.A.L. provided colorectal adenoma donor stool samples and G.D.W. and J.D.L. provided stool samples from Abx/PEG-treated human subjects. J.T.W. and S.O.M. contributed to visualization and writing. A.O., A.S.D., S.O.M., D.V.W., M.S., P.S.P., D.J.M., and J.T.W. contributed to writing, reviewing and editing with contributions from additional collaborators. A.O. and A.S.D. provided project administration and funding acquisition.

## METHODS

### RESOURCE AVAILABILITY

#### Lead Contact

Further information and requests for resources and reagents should be directed to and will be fulfilled by the Lead Contact.

### Materials Availability

All unique/stable reagents generated in this study are available from the Lead Contact through a completed Materials Transfer Agreement.

### Data and Code Availability

This study does not report original code or newly generated large-scale sequencing datasets. The previously published shotgun metagenomic sequencing data from the EDIA cohort analyzed in this study are available through the NCBI Sequence Read Archive under BioProject accession PRJNA821542. All other data generated in this study are provided in the article and Supplemental Information. Any additional information required to reanalyze the data reported in this paper is available from the Lead Contact upon request. This paper does not report original code.

## EXPERIMENTAL MODEL AND SUBJECT DETAILS

### Gnotobiotic mouse experiments

Mice were maintained in SPF or GF conditions when appropriate. C57BL/6 mice were purchased from Charles River. All experiments were performed on female mice between 8-12 weeks old. For gnotobiotic experiments, female mice were maintained in an Isocage system (Tecniplast). All animal procedures were approved by the Institutional Animal Care and Use Committee at Harvard Medical School.

8-12 weeks old C57BL/6 female mice were purchased from Charles River and maintained at the Harvard Center for Comparative Medicine facilities in Harvard Medical School. Mice were housed in specific pathogen-free conditions or GF conditions when appropriate. All mice were kept in a 12–12 h light–dark cycle, the main room temperature set at 20–26 °C and humidity of 40–65%. Mice had ad-libitum access to food and water. All mice were used following animal care guidelines from Harvard Medical School Standing Committee on Animals and the National Institutes of Health. Stool samples were approved for research use at Harvard Medical School under Non-Human Subjects research determination protocol IRB19-1887.

### Human Stool and Bile Collection

#### Stool collection from non-pregnant healthy humans from Boston

Samples were collected for a previously published study^98^ and banked samples were used in this study. Briefly, healthy human participants (14 females and 19 males, with replicate donations from 2 females and 4 males) with ages ranging from 23 to 38 were consented to participate under Committee on the Use of Humans as Experimental Subjects (COUHES) protocol number 1510271631 under the protocol “Culturing of Bacterial Strains within Healthy Individuals.” Age, sex, race and ethnicity, and diet of the subset of participants used are provided in (**Data Table 4**). The ancestry and socioeconomic status of these participants is not known. Although we do not anticipate that these factors would affect our results, the lack of information on these metrics is a limitation of the work. The participants gave their written informed consent to participate in this study. The above-referenced study was reviewed and approved by the Ethical Review Board of the Massachusetts Institute of Technology (MIT, Cambridge, MA), COUHES protocol number 1510271631. Stool samples were approved for research use at Harvard Medical School (HMS) (IRB19-1887).

#### Stool collection from pregnant and non-pregnant donors from Finland

Samples were collected as part of the previously published EDIA study^74,75^ and banked samples were used in this study. Frozen stool samples were collected from a longitudinal cohort of 26 pregnant participants at three timepoints (third trimester, at delivery, and ≥12 months postpartum; age range at delivery, 24.4–45.6 years), nine additional pregnant participants sampled during the third trimester, and 30 non-pregnant female controls. Participants were women of Finnish ancestry, and sample collection was approved by the Joint Municipal Authority of Tampere University Hospital, Tampere, Finland. Chronic disease, disease diagnosed during pregnancy, and antibiotic use are reported in **Data Table 5**. Socioeconomic status was not available. Stool samples were approved for research use at Harvard Medical School (IRB25-0818)

#### Stool collection from patients with a history of colorectal adenomas at Massachusetts General Hospital

Banked frozen stool samples collected from 38 male and 15 female patients with a history of colorectal adenomatous polyps (adenomas) at Massachusetts General Hospital were used in this study. Banked samples were collected from patients at a baseline flexible sigmoidoscopy procedure as part of a separate study. At this baseline study visit, blood pressure was measured prior to the flexible sigmoidoscopy procedure, and stool was collected during the procedure approximately 30-60 min later. All participants provided written informed consent for research use of their samples, and the collection protocol was approved by the Harvard Cancer Consortium Institutional Review Board (HCC IRB #19-402). Sex, BMI, age, blood pressure, and antihypertensive medication use are reported in Data Table 6.

#### Stool collection from Abx/PEG-treated donors from the University of Pennsylvania

Samples were collected for a previously published study,^67^ and banked samples were used in this study. Briefly, 10 healthy volunteers from the omnivore diet arm between the ages of 22-68 were included for analysis. Age, sex, race, and BMI can be found at (Tanes et al 2022).^67^ Key exclusion criteria included inflammatory bowel disease, celiac disease, or other chronic intestinal disorders; prior bowel resection surgery other than appendectomy; baseline bowel frequency less than every 2 days or greater than 3 times daily; creatinine concentration greater than the upper limit of normal; diabetes mellitus; currently smoking; body mass index (BMI) <18.5 or >35; and use of antibiotics or probiotics in the prior 6 months. On days 6, 7, and 8, inpatient participants received vancomycin 500mg orally every 6 h and neomycin 1000mg orally every 6 h. On day 7, participants consumed 4L of polyethylene glycol (PEG) based bowel purgative (GoLytely®). The University of Pennsylvania Institutional Review Board (IRB) approved the research protocol and considered it exempt from clinical trial registration requirements based on the protocol’s stated objectives. Stool samples were approved for research use at Harvard Medical School (HMS) (IRB26-0355).

#### Bile collection from primary-sclerosing cholangitis (PSC) patients

Samples were collected for a previously published study^50^ and banked samples were used in this study. Briefly, bile was collected prospectively during endoscopic retrograde cholangiopancreatographies (ERCPs) at Brigham & Women’s Hospital (BWH, Boston, MA) for patients with PSC. A sterile catheter was passed through the scope, and bile was carefully aspirated upon entry to minimize contamination. For patients without PSC (n=4), bile was obtained via gallbladder resection (n=1), via cholecystectomy (n=2), or received an ERCP due to biliary stricture (n=1). The age, sex, ancestry, race, or ethnicity, and socioeconomic status of these participants is not known. Although we do not anticipate that these factors would affect our results, the lack of information on these metrics is a limitation of the work. Bile samples were approved for research use at HMS (IRB19-1891).

#### Ethics

All mouse experiments were performed under the approval of the HMS Institutional Animal Care and Use Committee. Human stool collection was performed under the approval of the UPenn, MGH, and MIT Institutional Review Boards as well as The Joint Municipal Authority of Tampere University Hospital, Tampere, Finland. Human bile collection was performed under the approval of MGH and BWH Institutional Review Boards. Colon organoids used were obtained in accordance with the ethical protocols ‘Ethics Committee of Basel, EKBB, no. 2019-02118’ and ‘Comitato Etico Territoriale Lombardia 5, Autorizzazione n. 3631’, following written informed consent from the patients prior to enrollment.

## METHODS DETAILS

### Bacterial Culturing

All human gut bacteria were cultured in an anaerobic chamber (Coy Laboratory Products) with a gas mixture of 5% hydrogen and 20% carbon dioxide (balance N_2_) unless otherwise stated. Individual strains were grown in brain heart infusion (BHI) (Bacto, BHI) media supplemented with 1% Vitamin K1-Hemin Solution (Becton, Dickinson, 212354), 1% Trace Mineral Supplement (ATCC, MD-TMS), 1% Vitamin Supplement (ATCC, MD-VS), 5% heat inactivated Fetal Bovine Serum (Gibco, A5670701), 1 g/L cellubiose, 1 g/L maltose, 1g/L fructose (BHI+). Additionally, plates were supplemented with 0.1% arginine, and liquid media was supplemented with 0.5% arginine. *Escherichia coli* strains were grown aerobically at 37 °C in Luria-Bertani (LB) or Terrific Broth (TB) media supplemented with kanamycin (Kan) and chloramphenicol (CAM) to select for the pET28a(+) and pLysS plasmids. All experiments were performed in triplicate and repeated twice with similar results unless otherwise stated.

### UHPLC-MS analysis of steroid hormones

#### Unit selection

Steroids extracted from liquids (bacterial cultures, bile) are reported in μM. Steroids extracted from solids (feces) are reported as fmol/mg or pmol/mg wet mass.

#### Sample preparation for bacterial and pure protein samples

All samples were extracted as previously described.^50^ Briefly, 75 μL of water was added to 25 μL sample, followed by 100 μL of 1 μM ganaxolone (internal standard) in methanol. Sample and internal standard were incubated with 100 μL of ammonium acetate, pH 5.5, for 1 h before twice extracting with 300 μL of ethyl acetate:hexanes (80:20). Dried extracts were resuspended in 400 μL of LC-MS grade acetonitrile and diluted 1:10 in 1 μM pregnenolone-d2,13C2 (injection standard) in acetonitrile.

#### Sample preparation for human bile quantification

Extracts were prepared as above for bacterial and pure protein samples. For bile, 100 μL of sample was extracted after being digested with ß-glucuronidase/arylsulfatase (Sigma 10127698001) at 37 °C for 16 h.

#### Sample preparation for mouse feces quantification

Fecal pellets were homogenized in pre-weighed tubes in 600 μL of 80% methanol. Supernatants were extracted as described above.

### LC-MS for bacterial, mouse, and human sample extracts

Extracts were analyzed on a Orbitrap Exploris 120 Mass Spectrometer (ThermoFisher) equipped with a Vanquish Flex UHPLC (ThermoFisher) and separated using a Phenomenex Kinetex C18 column (1.7 μm particle size, 2.1x100). Flow rate was 0.45 mL/min with mobile phase A as water with 0.1% formic acid and mobile phase B as acetonitrile with 0.1% formic acid. The column temperature was maintained at 15 °C. Steroids were separated with the following gradient: 0-1 min: 5% B isocratic, 1-4 min: 5-30% B, 4-8 min: 30-44% B, 8-13 min: 44-99% B, 13-15 min: 99% B isocratic, 15-15.1 min: 99-5% B, 15.1-18 min: 5% B isocratic. For mass spectrometry, an APCI source was used. The ion transfer tube was maintained at 275 °C and the vaporizer temperature was maintained at 410°C. The masses of the THB isomers (loss of water, mass 315.2319 m/z), the 11βOH THP isomers (loss of water, mass 299.2369 m/z), 11oxo THP isomers (mass 315.2319 m/z), ganaxolone (internal standard, loss of water, mass 315.2682 m/z), and pregnenolone-d_2_,^13^C_2_ (internal standard, loss of water, mass 303.2562 m/z) were ionized in positive mode at 1.7 μA and 70% RF lens with 60k orbitrap resolution via single ion monitoring (SIM). For tissue and fecal sample analysis, a SecurityGuard ULTRA guard column (Phenomenex) was also used.

### Bacterial co-culturing experiments

Initial starter cultures were grown from single colonies in BHI+ media for 2-3 d. Experimental cultures were made by diluting monocultures 1:100 and co-inoculating the relevant strains in triplicate into 4 mL fresh liquid media. 100 μM of the corresponding substrate or vehicle control was supplemented into the media, and cultures were grown at 37°C for time course experiments. An aliquot of culture (0.2 mL) was collected and used for metabolite quantification.

### Screening *E. lenta* strains for 11-oxidation activity

Initial starter cultures were grown from single colonies in BHI+ media for 2-3 d. Experimental cultures were made by diluting starter cultures of *E. lenta* strains to a 1:100 dilution in 1.5 mL BHI+ media in a sterile deep 96-well plate. Substrate was added to co-cultures at a 100 μM concentration. Empty wells were filled with 1 mL of PBS. Plates were covered with an AeraSeal film (Sigma-Aldrich A9224) and incubated for 2 d at 37 °C. Samples were removed from the incubator for extraction and mass spectrometry analysis.

### Bioinformatics search for 11β-HSDH candidate gene

BLASTP searches were performed against the *E. lenta* type strain, *E. lenta* DSM 2243 (taxid:479437), on NIH National Center for Biotechnology Information (NIH NCBI) using the RefSeq Select proteins database. Six known *Eggerthella sp*. HSDH enzymes 3βHSDH2 (GenBank: ACV55294.1), 3αHSDH (GenBank: ACV54671.1), 3βHSDH (GenBank: ACV54349.1), 3βHSDH (GenBank: ACV54350.1), 3βHSDH (Genbank: ACV54351.1), and 3βHSDH1 (Genbank: ACV54192.1) were used as query sequences. Subsequently, the nucleotide sequences of all novel protein sequences producing significant alignments to the initial queries were used in BLASTn searches in the whole-genome shotgun contigs database limited by the BioProjectID 412637. Sequences that were present in strains with 11-oxidative activity were selected for cloning.

### Cloning putative 11βHSDH genes

Gene blocks from gene candidate 11βHSDH genes in *E. lenta* DSM 2243 (Elen_1987, Elen_2515, and Elen_2188) were synthesized by Twist Biosciences. The backbone was generated by linearizing a pET-28b vector containing an N-terminal His6 tag via PCR. PCR products were separated in 1% agarose gel electrophoresis and excised. Excised product was then purified using a Spin DNA Gel Extraction kit (Monarch, T1120L ). 11βHSDH gene blocks were joined to linearized pET-28b vector using Gibson assembly and full plasmids transformed into TOP10 competent cells. Transformed cells were grown in LB medium supplemented with Kan (50 μg/mL) at 37 °C overnight and plasmid prepped (ZR Plasmid Miniprep - Classic, Zymo Research, ZD4015). Full plasmid sequencing was performed at Plasmidsaurus prior to transformation into BL21 (DE3) pLysS *E. coli* (Novagen).

### Protein purification and activity assay of putative 11βHSDH enzymes

BL21 (DE3) pLysS *E. coli* plasmids were grown to saturation in 10 mL TB medium supplemented with Kan (50 μg/mL) and CAM (30μg/mL) at 37 °C. Starter cultures were used to inoculate 1 L of Kan/CAM TB media and expression of the N-terminal His6 fusion proteins was induced at an OD600 of 0.5-0.6 with 500 μM of isopropyl β-D-1-thiogalactopyranoside (IPTG) (Sigma-Aldrich, I5502-1G). Induced cells were incubated at 18 °C for 20 h then pelleted by centrifugation (15 min at 4,200 rpm at 4 °C). Afterwards, cell pellets were resuspended in 25 mL of ice-cold lysis/wash buffer (50mM HEPES, 30 mM imidazole, 250 mM NaCl, pH 8.0) and lysed by sonication. Cell debris was removed by centrifugation (20 min at 12,000 x g at 4 °C). Using a Nickel NTA column, protein was washed with lysis/wash buffer until no color change was detected using Bradford reagent on aliquots. Elution buffer (50mM HEPES, 250mM Imidazole, 250 mM NaCl, pH 8.0) was then used to elute the protein from the column. Afterwards, imidazole was removed from purified protein using dialysis buffer (50mM HEPES, 250mM NaCl, 10% glycerol, pH 8.0) and dialysis columns (Econ-Pac 10DG Desalting Columns, BIO-RAD). Protein expression was confirmed by western blot and SDS page analysis. For activity assay, 100 µM of DCA or **1** was added to purified protein. After incubation at 37 °C for 4 h, the reaction was quenched at -80 °C. A 25 µL aliquot was removed for UHPLC-MS analysis.

### Metagenomic analysis of the EDIA cohort

Shotgun metagenomic sequencing data from stool samples in the previously published EDIA cohort were analyzed as previously described.^74^ Briefly, sequencing data were quality controlled using KneadData v0.7.2, and taxonomic profiling was performed using MetaPhlAn v2.9.21.^99^ For the present study, genus-level taxonomic profiles from 25 pregnant and 23 non-pregnant samples were used to compare the prevalence and relative abundance of Eggerthella, Gordonibacter, Bacteroides, and Parabacteroides between groups. Differences in the prevalence of Eggerthella and Gordonibacter between pregnant and non-pregnant samples were assessed using Fisher’s exact test, and differences in relative abundance were assessed using unpaired Wilcoxon rank-sum tests.

### Cell and organoid culture

Human embryonic kidney-293 (HEK-293, female, ATCC-CRL-1573) cells and human SW-620 (male, ATCC-CCL-227) colon cells were obtained from the American Type Culture Collection (ATCC, Manassas, VA, USA). HEK-293 cells were stably transfected with a plasmid for hHSD11B2.^100^ The cells were cultured in Dulbecco’s modified Eagle medium (DMEM, BioConcept, Allschwil, Switzerland, 1-26F03-I, 4.5 g/L glucose and 4 mM L-glutamine), supplemented with heat-inactivated 10% fetal bovine serum (FBS, Southamerica, Biowest, Nuaillé, France, S1810-500), 100 U/mL penicillin, 0.1 mg/mL streptomycin (BioConcept, 4-01F00-H), 10 mM HEPES buffer (BioConcept, 5-31F00-H, pH 7.4), and 1% MEM non-essential amino acids (BioConcept, 5-13K00-H) referred to as cDMEM.

V79 (chinese hamster, male, ATCC-CCL-93) lung fibroblast cells were sub-cultivated in 10 cm^2^ dishes with DMEM (BioConcept, 1-26F03-I, 4.5 g/L glucose and 4 mM L-glutamine) supplemented with 100 U/mL penicillin, 0.1 mg/mL streptomycin, sodium pyruvate (Sigma-Aldrich, St. Louis, MO, USA, S8636, 1 mM) and 10% FBS. For treatment, DMEM (Sigma-Aldrich, D1145, containing 4.5 g/L glucose, but no phenol-red) was supplemented with 100 U/mL penicillin, 0.1 mg/mL streptomycin, sodium pyruvate (1 mM) and L-glutamine (BioConcept, 5-10K00-H, 4 mM) was used (from now on referred to as treatment media).

Cells were split when reaching 70-80% confluence. Cells were passaged using 1x Trypsin-ethylenediaminetetraacetic acid (EDTA) solution (Sigma-Aldrich, T4174). Cells were counted with the EVE™ Automatic cell counter (Witec AG, Sursee, Switzerland) and trypan blue was used to assess cell number and their viability. For experiments, cells were used at passage numbers 10-20.

Colon organoids were kindly provided by the group of Prof. Dr. Salvatore Piscuoglio. The organoids were obtained from healthy tissue and the procedure was optimized as described earlier.^101^ P3 is a sample obtained from a 51 year old male patient, P2 from a 60 years old female patient, and P1 from a 71 year old female patient. Organoids were sub-cultivated in 6-well plates using Corning® Matrigel® (Sigma-Aldrich, CLS356231-1EA) as extracellular matrix. For subculturing, culture media was removed and 600 µL *TryplE* (Thermo Fisher Scientific, Basel, Switzerland, 12605010) added per 6-well. Matrigel® domes were scratched with a cell scraper, transferred to a 15 mL tube, and the plate was washed with additional 600 µL *TryplE.* Organoids were then incubated for 14 min at 37°C in a water bath and, after 7 min, dissociated by pipetting. If Matrigel® remained, the procedure was repeated with additional 500 μL *TryplE* and incubation for 6 min at 37°C in a water bath. The tube was filled up with cold DMEM (Thermo Fisher Scientific, 11995-040) and centrifuged for 5 min at 300 x g at 4°C. Supernatant was removed, followed by resuspension in Matrigel® (10 drops of 20 µL Matrigel® per 6-well). After 20 min at 37°C and 5% CO_2_, 2 mL IntestiCult OGM media (# 06010, StemCell Technologies, Basel, Switzerland) was added. Cultures were maintained under standard conditions at 37°C in a humidified atmosphere with 5% CO_2_ and monthly tested for absence of mycoplasma.

### Pellet production from colon organoids

Colon organoids were washed with ice-cold phosphate-buffered saline (PBS) and Matrigel® drops were detached from the wells using a cell scraper. The resulting suspension was transferred into a 15 mL Falcon tube, with 2 mL of ice-cold PBS used per well. The tube was shaken gently and centrifuged at 200 x g for 5 min at 4°C. The PBS was aspirated as much as possible to the organoid-containing Matrigel® phase without disturbing the Matrigel® layer. The pellet was resuspended in ice-cold PBS (2 mL) and subjected to another round of centrifugation at 300 x g for 5 min at 4°C. This process of PBS aspiration, resuspension in fresh PBS, and centrifugation was repeated twice more, with centrifugation steps performed at 400 x g for 5 min at 4°C. To dissociate the Matrigel® matrix, Dispase II (Thermo Fisher Scientific, 17105041) (250 µL per well) was added to the pellet, and the mixture was gently pipetted. The suspension was incubated at 37°C for 40 min. Dispase II activity was inactivated by adding 5 mL of 0.5 M EDTA (Thermo Fisher Scientific, 15575020) and 7 mL of ice-cold PBS. The suspension was centrifuged at 400 x g for 5 min at 4°C and the resulting pellet was washed once with PBS (2 mL). After another centrifugation at 400 x g for 5 min, the pellet was resuspended in 1 mL of PBS and transferred to a 1.5 mL microcentrifuge tube. The sample was centrifuged again at 400 x g for 5 min, the supernatant was carefully removed, and the final pellet was snap-frozen on dry ice. One sample/6-well was prepared per patient-derived organoid.

Organoid characterization was performed by RT-qPCR and Western Blot. For further experiments, P1 was used, which showed the highest HSD11B2 protein expression on Western blots. Furthermore, lysate assay was performed to assess enzyme activity prior to continuing with intact organoid activity assay.

### Lysate HSD11B2 activity assay

HSD11B2 activity assay was performed as described earlier.^102^ Shortly, stably transfected hHSD11B2 HEK-293 cells, HEK-293 cells transiently transfected with mouse HSD11B2 (transient transfection was achieved with the calcium-phosphate precipitation method) and 1x pellet of (Specific Patient ID P1) colon organoids were lysed, prepared with the respective treatments and incubated with 10 nCi tritium-labeled cortisol, 40 nM cortisol, 500 µM NAD^+^ for 10 min (2 h for colon organoids) at 37°C. After incubation, the reactions were stopped by adding an excess of cortisol/cortisone in methanol (2 mM each), followed by separation of the steroids by thin-layer chromatography (TLC) and analysis of the conversion of cortisol to cortisone using a scintillation counter. Enzyme activity was calculated as cortisone / (cortisone + cortisol) ratio. Dimethylsulfoxide (DMSO, 0.1%) served as non-inhibiting negative control and 10 µM (GA) as inhibiting positive control. Half maximal inhibitory concentration (IC_50_) values were calculated relative to the 0.1% DMSO vehicle control. Mean and standard deviation (SD) were calculated from at least three independent experiments performed in technical duplicates. For the organoid assay, one patient-derived organoid (P1) was used.

### Intact cell HSD11B2 activity assay

HSD11B2 activity assay was performed as described earlier.^73,102^ Briefly, SW-620 cells were seeded in a density of 500’000 cells/well in a 12-well plate for 24 h. Cells were then washed 2x with serum-free media (culture media without 10% FBS) and prepared with the respective treatments and 10 nCi tritium-labeled cortisol and 40 nM cortisol in serum-free media (4h incubation). Supernatant was then collected and extraction performed by adding twice the amount of ethyl acetate and shaking 4 x 15 s at 1’800 rpm, followed by transferring the upper phase to a new tube and evaporating to dryness in a Genevac EZ-2 (SP Scientific, Warminster, PA, USA). Resuspension was done with 40 µL of cortisol/cortisone (each 2 mM). Cortisone and cortisol were separated by TLC, scraped, and analyzed by scintillation counting. Enzyme activity was determined as cortisone / (cortisone + cortisol) ratio. The assay was conducted in three independent experiments and technical duplicates.

### Colon organoid HSD11B2 activity assay

For each experiment 1 x 6-well was used. Organoids were dissociated as described in the cell and organoid culture section. After centrifugation the pellet was resuspended in 700 µL Advanced DMEM (Thermo Fisher Scientific, 12634010) supplemented with 1x glutaMAX (Thermo Fisher Scientific, 35050061), 1 mM HEPES (BioConcept, 5-31F00-H), 50 ng/µL EGF (Thermo Fisher Scientific, AF-100-15-100UG), 100 ng/mL Noggin (Thermo Fisher Scientific, 120-10C-50UG), 10 nM Gastrin (Sigma-Aldrich, G9145-0.1MG), 500 nM A83-01 (StemCell Technologies, 72024), 10 µM Y27632 (StemCell Technologies, 72304), 5 µM DAPT (StemCell Technologies, 72082), 1 mM N-Acetylsysteine (Chemie-Brunschwig, Basel, Switzerland, FLUF003631-25) and 1x B27 (Thermo Fisher Scientific, 12587010).^103^ 70 µL were seeded into Corning™ Costar™ 96-Well, Ultra-Low Binding, Flat-Bottom Microplate (Fisher Scientific, Thermo Fisher Scientific, 10554961) wells. Organoids were cultured in the supplemented advanced DMEM for six days with a media change after the first two days (added 100 µL fresh media, centrifuged plate at 300 x g for 3 min, replaced with 100 µL fresh media). Organoids were then treated for 6 h with the respective inhibitor and with 10 nCi tritium-labeled cortisol and 40 nM cortisol. The reaction was stopped by adding 10 µL of stop mix (2 mM each cortisol/cortisone). 10 µL was separated by TLC and conversion of cortisol to cortisone was measured by scintillation counting. Enzyme activity was calculated as cortisone / (cortisone + cortisol) ratio. The assay was performed in three independent experiments from one patient-derived organoid each in technical duplicates.

### Transactivation assay

V79 cells (150’000 cells/well) were seeded in 24-well plates. After 3 h of incubation, cells were transfected with pMMTV-lacZ β-galactosidase (MMTV-LacZ) reporter (0.5 ng/μL), pCMV-LUC luciferase (cluc) transfection control (0.1 ng/µL),^104^ hMR (gift from Dr. Nourdine Faresse)^105^ and empty pcDNA3.1 (V79020, Invitrogen, Basel, Switzerland), or hHSD11B2 cloned in a pcDNA3.1 backbone (0.5 ng/µL) with polyethylenimine (PEI) at a 1:3 DNA:PEI ratio in Opti-MEM. After 5 h, Opti-MEM was replaced with cDMEM, followed by incubation for 16 h to enable expression of the proteins. Cells were then washed once with PBS and cultured in treatment media (serum-free and phenol-red free) for 1 h at 37°C. The treatment medium was replaced with fresh treatment media containing the respective compounds. If two compounds were used in co-treatments, the antagonist was added for 1 h and then the agonist was added. After 24 h the cells were washed with PBS and lysed in 60 µL lysis buffer from the Tropix kit containing 1 µM DTT (Applied Biosystems, Foster City, CA), and frozen at -80°C for at least 20 min. Lysates (20 µL each) were used for β-galactosidase activity with the Tropix lysis kit. The accelerator was automatically injected (250 μL/s). For luciferase activity measurements, 20 µL of the lysates were used and 100 µL D-luciferin substrate solution (0.56 mM D-luciferin, 63 mM ATP, 0.32 mM CoA, 0.16 mM EDTA, 39.9 mM dithiothreitol, 9.6 mM MgSO_4_, 24 mM tricine (pH 7.8)) was automatically injected (250 μL/s). Luciferase and β-galactosidase activity were measured using a Cytation 5 reader (BioTek, Winooski, VT, USA). For half maximal effective concentration (EC_50_) curves, the values were calculated to 0.1% DMSO control. Mean and SD were calculated from three independent experiments, each performed in technical duplicates.

### Cell viability assay

For all cell-based intact assays, a cell viability assay was performed as described previously.^106,107^ Shortly, the highest concentrations used in the intact cell assays were tested for cytotoxicity in an assay using2,3-bis/2-methoxy-4-nitro-5-sulfophenyl)-2H-tetrazolium-5-carboxanilide (XTT). Briefly, 150 μL, 75 μL or 12.5 μL of a XTT/phenazine methosulfate (PMS) solution (7.5 μg/mL) for 12-well, 24-well or 96-well format, respectively, were added and absorbance at 450 nm and 650 nm (background) was measured 2 h later, 50% DMSO was used as a positive control. Results were normalized to the 0.1% DMSO control and mean and SD were calculated from three independent experiments performed in technical duplicates.

### Assessment of HSD11B2-mediated substrate conversion using UHPLC/MS-MS

For the evaluation of HSD11B2 enzyme activity in intact SW-620 cells substrate and product concentrations, *i.e.* cortisol (4-pregnen-11β,17,21-triol-3,20-dione; CAS 50-23-7) and cortisone (4-pregnen-17,21-diol-3,11,20-trione; CAS 53-06-5), or 3β,11β-diOH-5α-THP (5α-pregnane-3α,11β-diol-20-one; CAS: 565-89-9), 3α,11β-diOH-5α-THP (5α-pregnane-3β,11β-diol-20-one; CAS: 23930-29-2) and 3β-OH-11oxo-5α-THP (5α-pregnane-3β-ol-11,20-dione; CAS: 600-59-9); were determined as following.

SW-620 cells (250’000 cells/well) were seeded in a 24-well plate and incubated for 24 h. Cells were then treated with the respective concentrations indicated for different time points (t0 = 0 min, t24 = 24 h). Cell culture supernatant was taken and stored at -20°C. The experiment was performed in three independent experiments performed in two technical replicates, whereas cortisol treatments were performed in three biological replicates and one technical replicate.

Steroid concentrations were analyzed against the volume-matched calibrators prepared in the SW-620 serum-free cell culture media by serial dilution (1:2) within the range of 1000-15.62 nM. Calibrators were subsequently treated as samples. For solid-phase extraction (SPE), each sample (450 µL) was mixed with protein precipitation solution (100 µL, 0.8 M zinc sulfate in water/methanol; 50/50, v/v) containing (0.2 µg/mL) deuterium-labeled cortisol-D4 (4-pregnen-11β, 17, 21-triol-3, 20-dione-9,11,12,12-d4, CAS: 73565-87-4), as internal standard. Prior to SPE all samples were diluted with water (400 µL) and incubated in a thermoshaker (10 min at 4°C, 1350 rpm). Samples were centrifuged for 10 min at 18,100 x g and supernatants (900 µL) were transferred to Oasis HBL SPE (3 cc) cartridges (Waters, Milford, MA, USA), pre-conditioned with methanol and water (3 mL each). Samples were washed with water (3 x 1 mL) and methanol/water (3 x 1 mL, 10/90, v/v). Cartridges were allowed to dry for 5 min. Steroids were eluted with methanol (2 x 750 µL), evaporated to dryness (3 h, 35 °C), and reconstituted in methanol (50 µL) by shaking (20 min at 4°C, 1350 rpm), followed by sonication for 10 minutes. Samples were centrifuged for 10 min at 18,100 x g and supernatants were transferred into auto-sampler vials equipped with low volume inserts. Analytes were measured by UHPLC-MS/MS using an Agilent 1290 Infinity II UHPLC coupled to an Agilent 6495D triple quadrupole mass spectrometer equipped with a jet-stream electrospray ionization interface (Agilent Technologies, Santa Clara, CA, USA). Analyte (Injection volume 3 µL) separation at 40 °C was achieved within 9.0 min using a reverse-phase column (1.7 µm, 2.1 mm x 150 mm; Acquity UPLC BEH C18; Waters). The mobile phase consists of methanol (B: 100%) and methanol water (A: 5/95% v/v). The following gradient (B%) conditions were applied for elution at constant flow (0.3 mL/min): 0-10 min (50%-72%), followed by a column washout 10-12.5 min (72-100%). The column was post-run reconstituted to initial B% within 1.5 min prior further injections. Analyte-specific parameters are found in SI content Data Table 1 Data acquisition (Version 12.1, Update 3 Build 12.1.186 Agilent Technologies) and qualitative and quantitative analysis (Version B.10.0. Build 10.0.27, RRID: SCR_015040, Agilent Technologies) were performed by MassHunter.

### Cofolded model and docking

Because an experimentally determined structure of HSD11B2 was unavailable for these analyses, structural models were generated to investigate the inhibitory mechanism in both orthologues. Protein models were predicted using the cofolding methods Boltz-2 (version 2.1) and Chai Discovery (version Chai-1), with the primary FASTA sequence of human or mouse HSD11B2 as input together with the SMILES representation of the cofactor NAD⁺ and the respective natural substrate (cortisol for hHSD11B2 and corticosterone for mHSD11B2).

The models generated by Chai and Boltz exhibited highly similar overall folds and high confidence throughout most of the protein (predicted Local Distance Difference Test, pLDDT, between 70 and 100). The only notable difference was the C-terminal region of hHSD11B2, which showed lower confidence (pLDDT 50–70). The mHSD11B2 does not contain this extended C-terminal region. To further assess model reliability, both structures were compared with a homology model generated using SWISS-MODEL (https://swissmodel.expasy.org/) based on the closest available short-chain dehydrogenase/reductase (SDR) template. The overall fold and architecture of the substrate-binding pocket were highly consistent across all three models. The Chai model was selected for subsequent studies because it predicted the complete protein sequence and generated a more realistic three-dimensional conformation of the steroid substrate compared to Boltz, resulting in improved positioning relative to the NAD⁺ cofactor within the binding pocket.

Protein preparation and molecular docking were performed in Maestro (version 2024-3, Schrödinger, LLC, New York, NY, USA). The predicted protein-ligand complexes were prepared using the Protein Preparation Wizard with the OPLS_2005 force field. Structures were optimized and energy-minimized while restraining heavy atoms (RMSD cutoff 0.5 Å). Rotamers of key binding-site residues were manually adjusted where necessary to optimize the hydrogen-bonding network between the protein, cofactor, and substrate. Receptor grids were generated with Glide, allowing rotation of hydroxyl groups of serine and tyrosine residues within the binding pocket.

The natural substrates (cortisol or corticosterone) and the investigated progesterone metabolites (3β-OH/3α-OH, 11β-OH/11-oxo, and 5β-H/5α-H) were generated from SMILES strings using LigPrep at pH 7.4 +/- 0.5. The natural substrates were first redocked to validate the predicted binding mode, after which the progesterone metabolites were docked using Glide in extra precision (XP) mode. Following docking, all protein-ligand complexes were subjected to a final restrained optimization and energy minimization using the Protein Preparation Wizard. For mHSD11B2, the metabolite 3β5α11βOH-THP could not be placed in a plausible binding mode by Glide and was therefore positioned manually by superimposing its steroid scaffold onto that of the natural substrate.

### mRNA expression

SW-620 cells (500,000 cells/well) were seeded in a 12-well plate and incubated for 24 h. Cells were washed twice with PBS and RNA was extracted as described by the manufacturer with the QIAgen RNeasy Mini Kit (Applied Biosystems, Venlo, The Netherlands, 74106). For colon organoids, one pellet was used and processed with the QIAgen RNeasy Micro Kit (74004). NanoDrop One© (Witec AG, Sursee, Switzerland) was used to quantify RNA and reverse transcription was done with the Takara PrimeScript RT reagent kit (Takara Bio, Inc. Kusatsu, Japan, RR037A). RT-qPCR was performed on a QuantStudio 6 Pro System (Thermo Fisher Scientific). Oligonucleotide primers for RT-qPCR were purchased from Microsynth (Balgach, Switzerland) and sequences are found in SI content Data Table 3. RT-qPCR was performed with 4 ng of cDNA as a template and 200 nM of oligonucleotide primers with 1x KAPA SYBR FAST qPCR Master Mix, (Sigma-Aldrich, KK4618). Runs were started at 95°C for 5 min, followed by 40 cycles (10 s at 95°C, 15 s at 60 °C and 20 s at 72°C), and completed with a melting curve (95°C for 15 s, 60°C for 60 s and 95°C for 15 s). Ct-values were normalized to the housekeeping gene peptidylprolyl isomerase A (PPIA) (2−(Ct gene of interest−Ct PPIA)).

### Expression of HSD11B2 protein in patient derived organoids

One pellet of colon organoids was resuspended in RIPA buffer (Sigma-Aldrich, R8758) containing proteinase inhibitor cocktail (Roche, Basel, Switzerland, 11836153001). The protein concentration was measured with Pierce™ BCA Protein Assay Kits (Thermo Fisher Scientific, 23225) and samples were denaturated by adding with 4x Laemmli buffer (5 mM Trizma-HCl, pH 6.8, 10% glycerol [v/v], 0.2% sodium dodecyl sulfate [w/v], 1% bromophenol blue [w/v], substituted with 20% β-mercaptoethanol [v/v]) and boiling at 95°C for 5 min. Samples were stored at -20°C. 15 µg protein were loaded per sample. To separate proteins, 10% SDS-PAGE was run. To determine protein size, 5 µL of Precision Plus Protein™ All Blue Prestained Protein Standards (Bio-Rad Laboratories, #1610373, Cressier, Switzerland) was loaded on each gel. PVDF membranes (Merck KGaA, Darmstadt, Germany, IPVH00010) were blocked in 10% milk TBS-T (1.11 g/L Tween 20, 137 mM NaCl, and 20 mM Trizma Base, pH 7.6) at room temperature. Primary antibodies were incubated in 5% milk in TBS-T overnight at 4°C and secondary antibodies for 1 h in 5% TBS-T at RT. Used were the primary antibodies, Lamin B1 (Abcam, LUCERNA-CHEM, Lucerne, Switzerland ab133741, RRID: AB_2616597, 1:1000) HSD11B2 (Invitrogen, Thermo Fisher Scientific, PA5-59904, RRID: AB_2642555, 1:1000) and the secondary antibody raised in rabbit (Cell Signaling Technologies, Lane Danvers, MA, USA, 7074S, RRID: AB_2099233, 1:1000 for HSD11B2 and 1:2000 for Lamin B1). Immobilon Western horseradish peroxidase (HRP) substrate kit (Merck KGaA, Darmstadt, Germany, WBKLS0500) was used to visualize antibody-labeled proteins in a Fusion FX (Vilber Collégien, France). Fusion FX-associated FusionCapt Advance Software was used to assess relative protein expression by densitometry.

### Statistics

Statistical analyses were performed with GraphPad Prism Software 10.2 (RRID: SCR_002798, San Diego, CA, USA). Tests used are specified in the corresponding figure legend. p ≤ 0.05 was considered statistically significant (respective p-values are indicated). In instances where individual steroids were undetectable by UHPLC-MS, the estimated limits of detection (LODs) for each steroid were used for statistical tests (0.607 nM for (3α5α/3β5β)11βOH-THP, 0.267 nM for 3α5β11βOH-THP, 0.694 nM for 3β5α11βOH-THP, 0.311 nM (3α5α/3β5β)11keto-THP, 0.238 nM 3α5β11keto-THP, 0.229 nM 3β5α11keto-THP, 0.211 nM progesterone, 6.05 nM estradiol, and 7.35 nM estrone). LODs were estimated from both experimental and statistical experiments described by the Official Journal of the European Communities, Commission Decision of 12 August 2002; Implementing Council Directive 96/23/EC.^108^ Displayed are mean ± SD for cell and lysate experiments and mean ± SEM values for tissue and fecal experiments.

During the preparation of this work, the authors used ChatGPT (OpenAI) for editing and consistency checking across the main text and supplemental information. The authors reviewed and edited the output as needed and take full responsibility for the content of the published article.

