## Supplemental Figures and Tables for "Human Gut Bacteria Convert Endogenous Steroids into Host Cortisol Shuttle Inhibitors"

### SUPPLEMENTARY INFORMATION

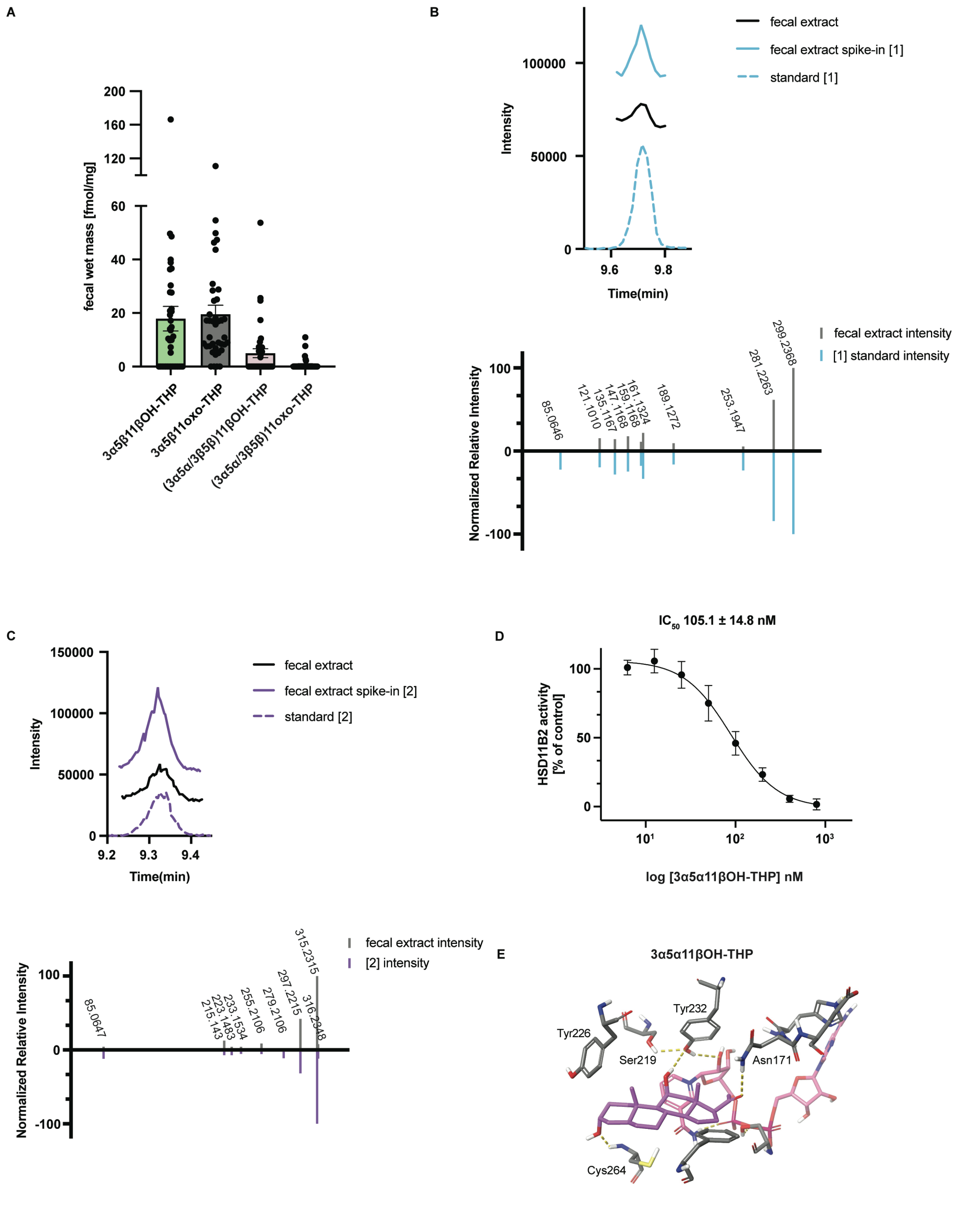

**Supplementary Figure S1. Identification and mass spectrometry-based validation of GALF metabolites in human feces**

(A) Concentrations of additional 11-oxygenated tetrahydroprogesterone (THP) isomers in feces from healthy human donors. Steroids were quantified by UHPLC-MS and are reported as fmol/mg wet mass (approximately nM) (n = 39 samples). Bars represent mean ± SEM.

(B) MS/MS-based validation of **1** in human fecal extracts. Top, extracted-ion chromatogram (EIC) for **1** in fecal extract and after spike-in with authentic **1**. Bottom, mirror plot showing comparison of normalized fragment intensities in authentic standard and fecal extract.

(C) MS/MS-based validation of **2** in human fecal extracts. Top, EIC for **2** in fecal extract and after spike-in with authentic **2**. Bottom, mirror plot showing comparison of normalized fragment intensities in authentic standard and fecal extract.

(D) Concentration-response curve for inhibition of human HSD11B2 by the known GALF 3α5α11βOH-THP in HEK293 cell lysates stably overexpressing hHSD11B2. Lysates were incubated for 10 min at 37°C with 40 nM cortisol, 10 nCi [$^3$H]cortisol, 500 μM $NAD^+$, and the indicated concentrations of 3α5α11βOH-THP. HSD11B2 activity (cortisone/[cortisol + cortisone]) was normalized to 0.1% DMSO (n = ≥3 independent experiments each in technical duplicate; mean ± SD). $IC_{50}$ was determined by nonlinear regression.

(E) Cofolded model of hHSD11B2 with docked 3α5α11βOH-THP showing the planar 5α-reduced A/B ring conformation and predicted hydrogen bonds between ligand and key active-site residues (e.g., CYS264, TYR232, ASN171).

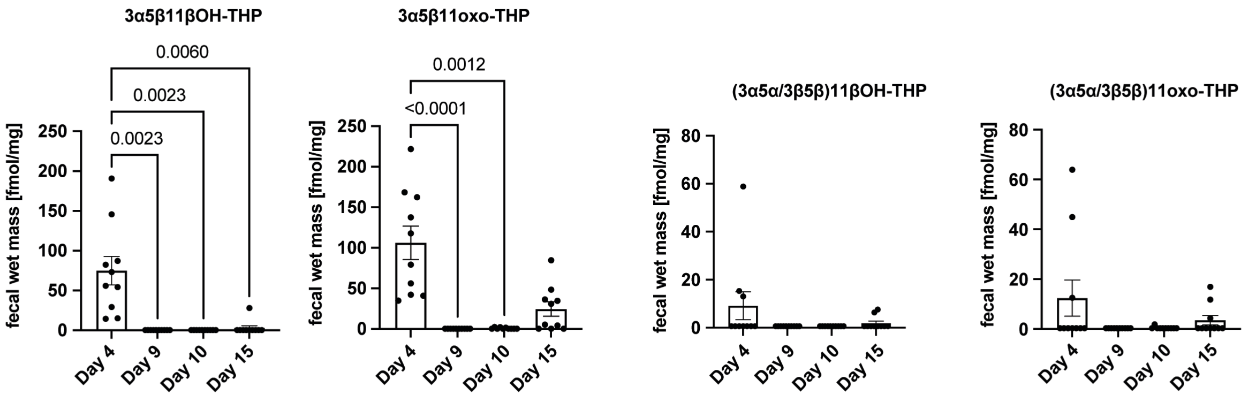


**Supplementary Figure S2. Microbiota depletion reduces C11-oxidized metabolites in human feces**

Fecal concentrations of 11-oxygenated THP derivatives in the longitudinal controlled-feeding/antibiotic (Abx)/polyethylene glycol (PEG) microbiota purge study.[1] Stool samples were collected on day 4 of the diet phase, immediately following the Abx/PEG phase (day 9), and during early recovery (day 10), and during late recovery (day 15). Metabolites quantified by UHPLC-MS include 3α5β11βOH-THP, 3α5β11oxo-THP, (3α5α/3β5β)11βOH-THP, and (3α5α/3β5β)11oxo-THP. Values are reported as fmol/mg wet fecal mass (approximately nM). Bars represent mean ± SEM (n=10 as in the omnivore group in Tanes et al., 2021 for each time point). Several C11-oxidized species, including the progestins 3α5β11βOH-THP and 3α5β11oxo-THP, were significantly depleted following the Abx/PEG phase. Statistical significance was assessed using a Friedman test followed by Dunn's multiple-comparisons test. Exact p values are indicated on the plots.

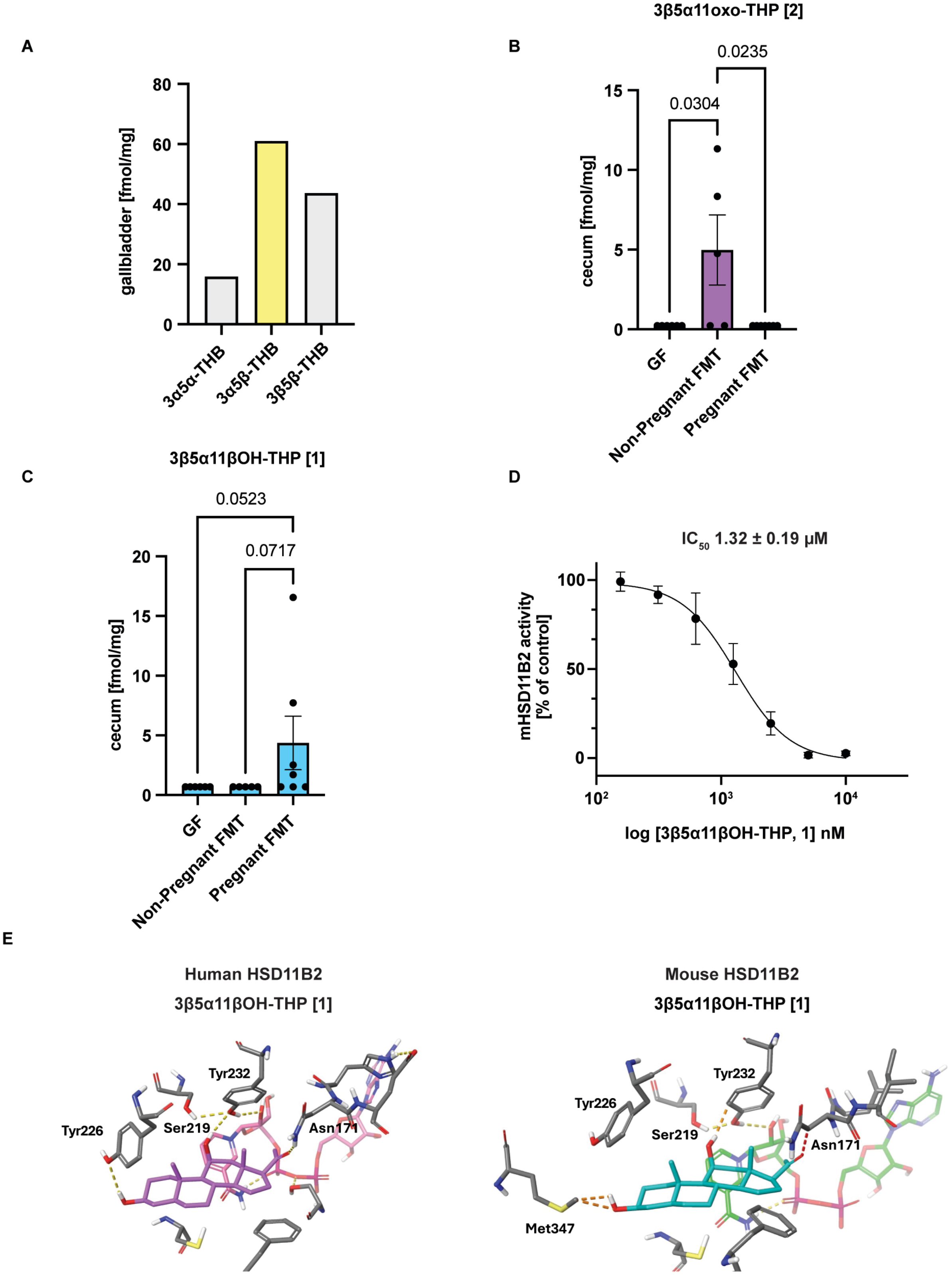

A
gallbladder [fmol/mg]
3α5α-THB
3α5β-THB
3β5β-THB
B
3β5α11oxo-THP [2]
cecum [fmol/mg]
0.0235
0.0304
GF
Non-Pregnant FMT
Pregnant FMT
C
3β5α11βOH-THP [1]
cecum [fmol/mg]
0.0523
0.0717
GF
Non-Pregnant FMT
Pregnant FMT
D
IC50 1.32 ± 0.19 μM
mHSD11B2 activity [% of control]
log [3β5α11βOH-THP, 1] nM
E
Human HSD11B2
3β5α11βOH-THP [1]
Tyr232
Tyr226
Ser219
Asn171
Mouse HSD11B2
3β5α11βOH-THP [1]
Tyr232
Tyr226
Ser219
Asn171
Met347

**Supplementary Figure S3. Levels and activity of GALF metabolites differ between mice and humans**

(A) Concentrations of 3α5βTHB and other THB isomers in mouse gallbladder bile, quantified by UHPLC-MS (due to limited bile volume in mouse gallbladders, each bar represents analysis of 7 pooled mouse gallbladders, so no error bars are shown).

(B-C) Cecal concentrations of **2** (B) and **1** (C) in germ-free (GF) mice, GF mice receiving fecal microbiota transplants (FMTs) from non-pregnant human donors, and GF mice receiving FMTs from pregnant donors. Steroids were quantified by UHPLC-MS and are reported as fmol/mg wet mass (n = 5-7 mice per group). Bars represent mean ± SEM. Kruskal-Wallis test followed by Dunn’s multiple-comparisons test was used to assess differences among groups.

(D) Concentration-response curve for inhibition of mouse HSD11B2 (mHSD11B2) by **1** in HEK293 cell lysates transiently overexpressing mHSD11B2. Assay conditions were as in Figure S1D. HSD11B2 activity was normalized to 0.1% DMSO (n = ≥3 independent experiments each in technical duplicate; mean ± SD). $IC_{50}$ was determined by nonlinear regression.

(E) Cofolded models of human and mouse HSD11B2 with docked **1**. Predicted hydrogen-bonding networks and side-chain environments around the 3β-OH, 11β-OH, and C20-oxo groups are shown.

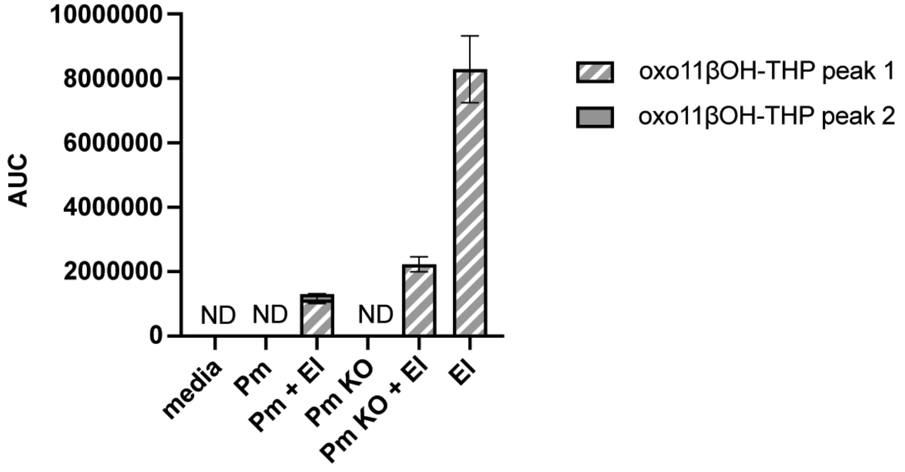


**Supplementary Figure S4 (related to Fig 3C). AUCs of unknown C11-oxygenated compounds produced by *E. lenta* 14A in mono- or co-culture in the presence of 3α5β11βOH-THP.**

*E. lenta* 14A (El) monocultures or co-cultures with *P. merdae* (Pm) or *P. merdae* Δ04016-18 (Pm KO) produced oxidized products with m/z values consistent with the intermediate 3oxo-5β11βOH-THP (m/z 315.2315). *P. merdae or P. merdae KO* monocultures alone were unable to produce these oxidized compounds. (Peak 1, RT: 9.66; Peak 2, RT: 9.82)

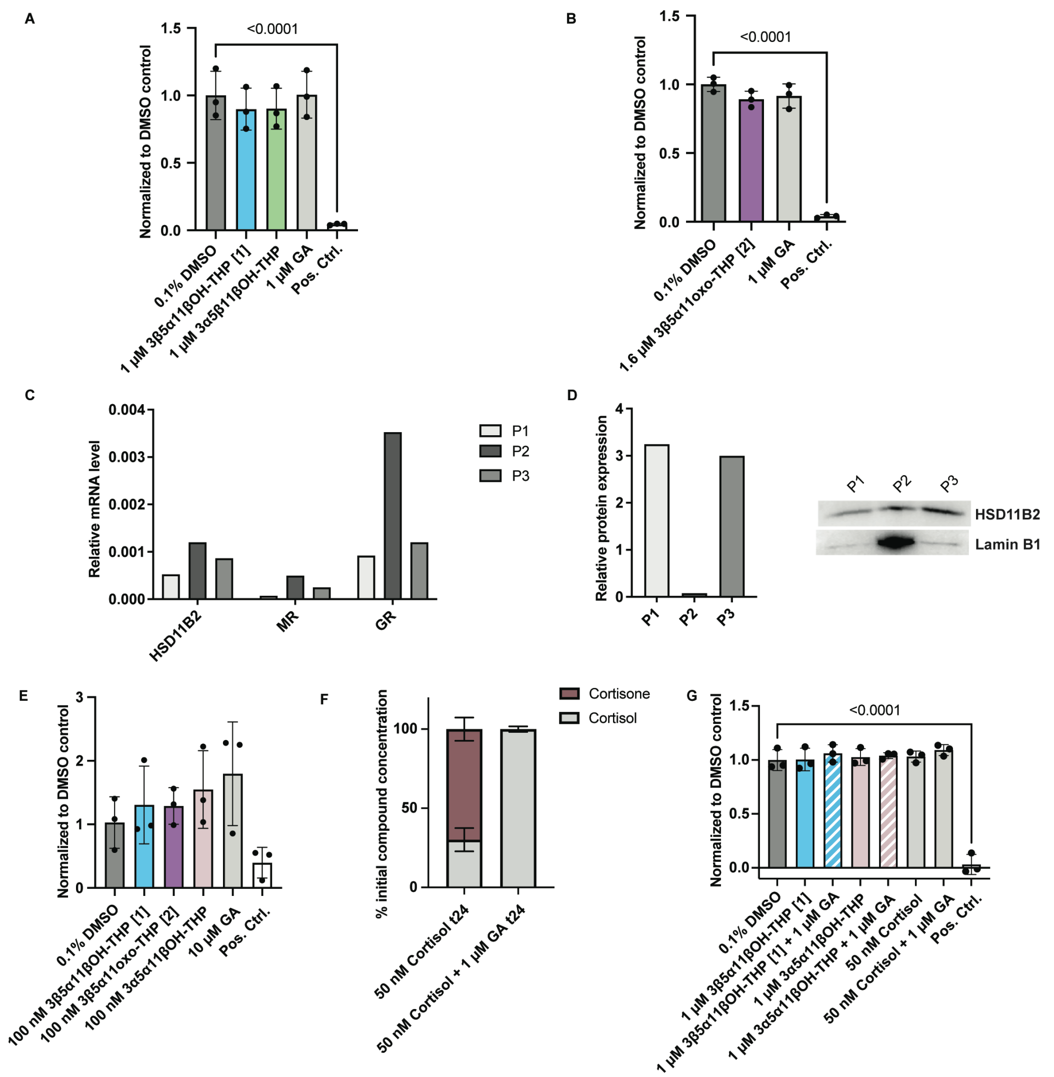


**Supplementary Figure S5. Cytotoxicity controls, organoid characterization, cortisol metabolism, and MR/GR specificity**

(A-B) XTT cell-viability assays in the SW-620 intact cell activity HSD11B2 experiments. Cells were treated for 4h and at the highest concentrations used in Fig. 4A-C under the conditions used in Fig. 4A-C (40 nM cortisol, 10 nCi [$^3$H]cortisol,with 3α5β11βOH-THP, **1**, **2**, GA or 0.1% DMSO). XTT/ phenazine methosulfate solution (PMS) was added for 2 h. As positive control, 50% DMSO was used. 0.1% DMSO served as a negative control. Data are normalized to 0.1% DMSO (n = 3 independent experiments in technical duplicate; mean ± SD).

(C) Relative mRNA expression of HSD11B2, MR, and GR in three patient-derived colon organoid lines (P1, P2, P3) measured by RT-qPCR. Expression is normalized to peptidylprolyl isomerase A (PPIA) (n = 1 organoid passage per donor, assayed in technical replicate).

(D) Protein expression of HSD11B2 in colon organoids determined by Western blot. Lamin B1 serves as a loading control. Representative blots and densitometry from n = 1 experiment are shown.

(E) XTT viability in colon organoids treated with **1** (100 nM), **2** (100 nM), 3α5α11βOH-THP (100 nM), GA (10 μM) for 6 h, 50% DMSO (or positive control) and 0.1%DMSO, as in the intact organoid HSD11B2 assay. Data are normalized to 0.1% DMSO (n = 3 independent experiments; mean ± SD).

(F) Cortisol metabolism in intact SW-620 cells. Cells were incubated with 50 nM cortisol ± 1 μM GA for 24 h, and cortisol and cortisone levels were quantified by UPLC-MS/MS. Bars represent percent of initial cortisol and cortisone (n = 3 independent experiments; mean ± SD).

(G) XTT viability in SW-620 cells under the conditions used in S5F and Fig. 4E with 3α5α11βOH-THP, **1**, cortisol, GA, or 0.1% DMSO. As positive control, 50% DMSO was used. 0.1% DMSO served as a negative control. Data are normalized to 0.1% DMSO (n = 3 independent experiments; mean ± SD).

For all panels, statistical analyses were performed using one-way ANOVA with Dunnett's multiple-comparison test unless otherwise stated; p values are indicated on the plots.

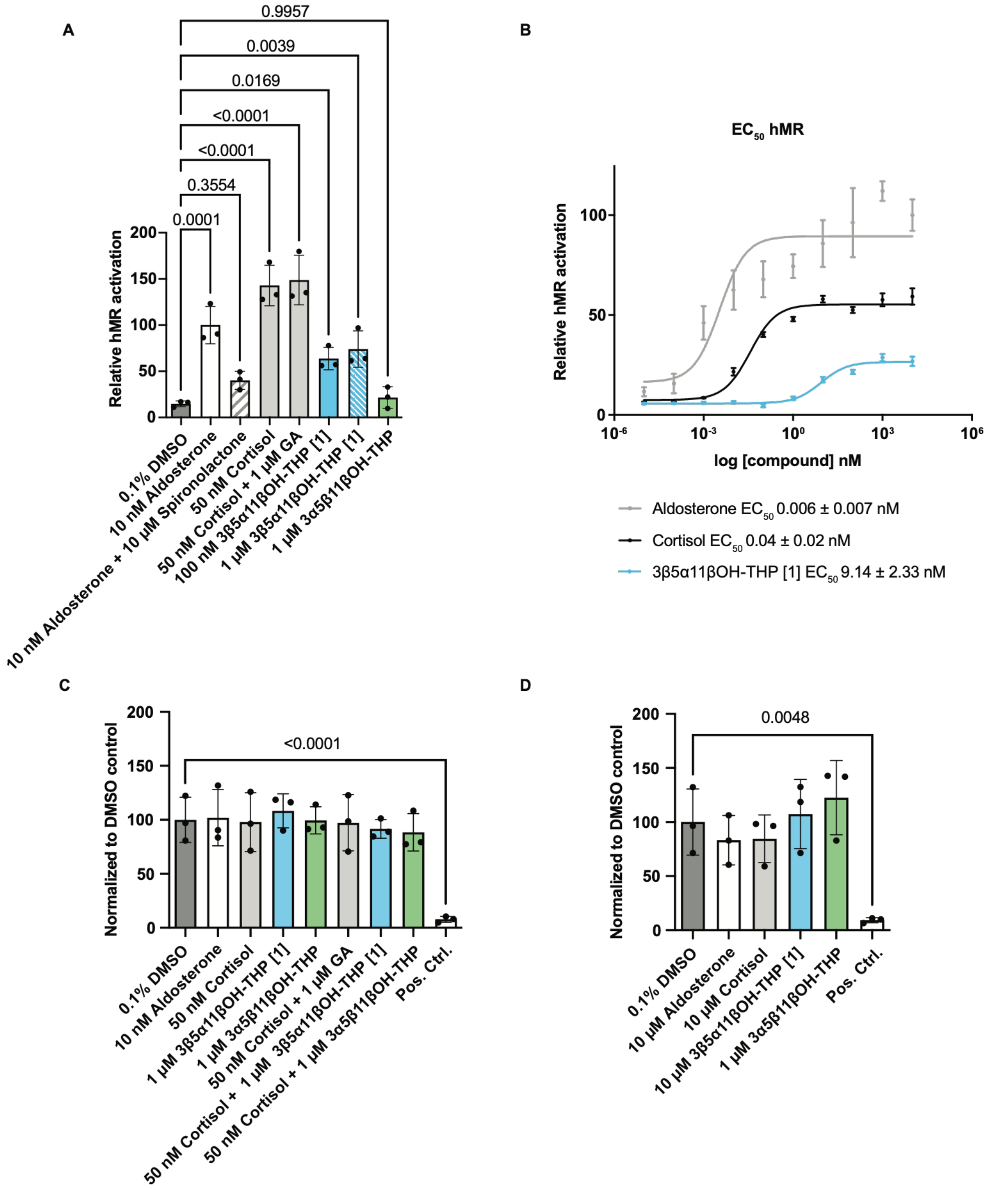


**Supplementary Figure S6. Nuclear receptor transactivation and specificity controls.**

(A) hMR activation in V79 cells transiently overexpressing hMR, MMTV-LacZ, and cluc, and empty pcDNA3.1. Cells were treated for 24 h with 10 nM aldosterone, 50 nM cortisol, 100 nM or 1 µM of compound **1**, 1 µM 3α5β11βOH-THP, or 10 µM GA as indicated. Where indicated, spironolactone or GA was added 1 h prior to agonist. MR activity was normalized to cluc and expressed relative to 10 nM aldosterone (n = ≥3 independent experiments in technical duplicate; mean ± SD).

(B) Concentration-response curves for hMR activation by aldosterone, cortisol, and **1** in the absence of HSD11B2. $EC_{50}$ values were obtained by nonlinear regression (n = ≥3 independent experiments; mean ± SD).

(C) XTT viability in V79 cells transiently overexpressing hMR and HSD11B2, MMTV-LacZ, and cluc. Cells were treated for the duration and at the highest concentrations used in (Fig 4F), Aldosterone, cortisol, **1**, 3α5β11βOH-THP, GA, or 0.1% DMSO. As positive control, 50% DMSO was used. 0.1% DMSO served as a negative control. Experiment performed as in Supp Fig 5A-B. Data are normalized to 0.1% DMSO (n = 3 independent experiments; mean ± SD).

(D) XTT viability in V79 transiently overexpressing hMR. Cells were treated for the duration and at the highest concentrations used in (A), Aldosterone, cortisol, **1**, 3α5β11βOH-THP, or 0.1% DMSO. Experiment performed as in Supp Fig 5A-B. Data are normalized to 0.1% DMSO (n = 3 independent experiments; mean ± SD).

For all panels, statistical analyses were performed using one-way ANOVA with Dunnett's multiple-comparison test unless otherwise stated; p values are indicated on the plots

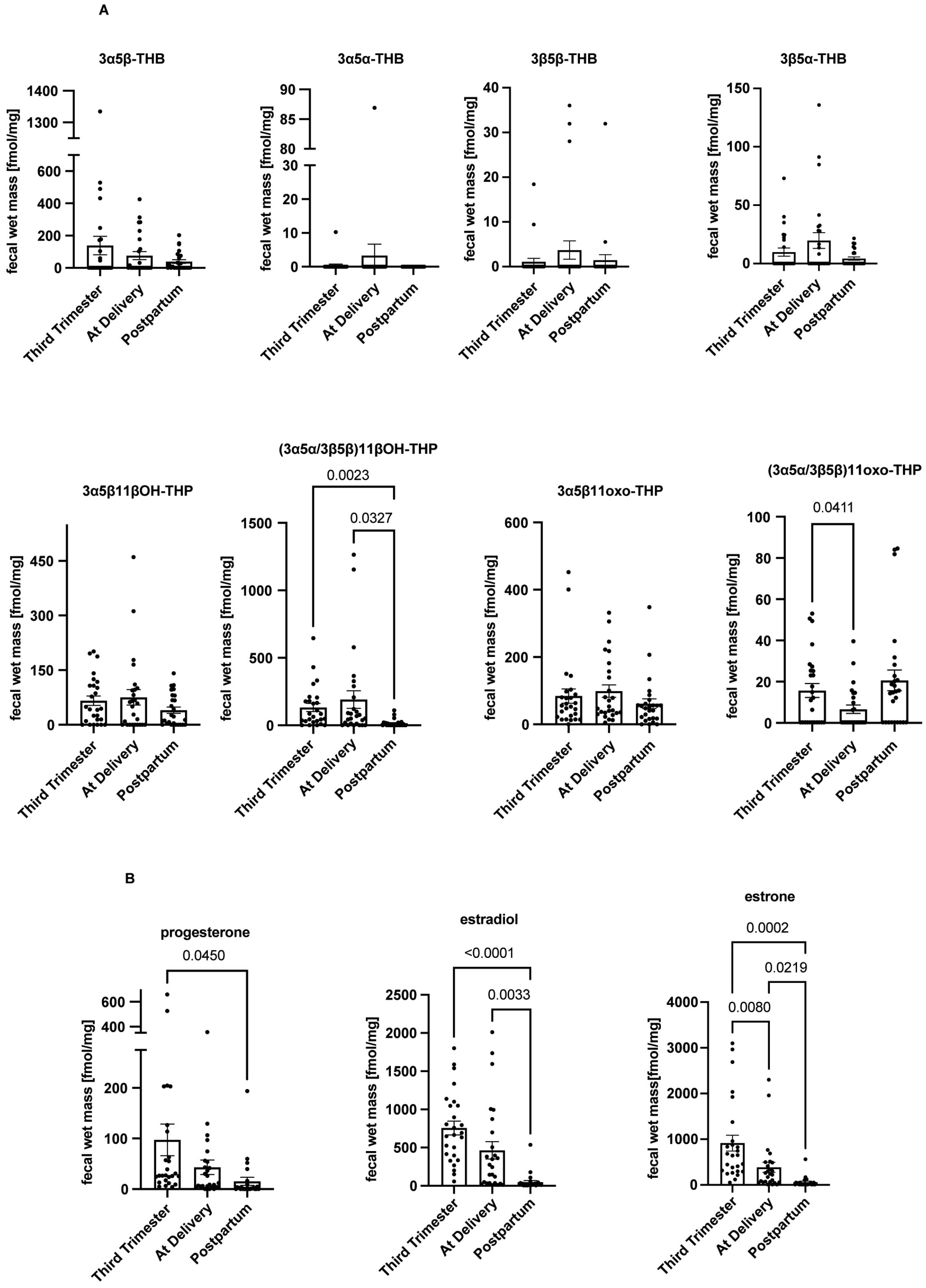


**Supplementary Figure S7. Extended fecal steroid profiling across pregnancy and postpartum**

(A) Fecal concentrations of 11-oxygenated corticoids and THP derivatives across pregnancy in the Finnish cohort. Steroids include 3α5β-THB, 3α5α-THB, 3β5β-THB, 3β5α-THB, 3α5β11βOH-THP, 3α5α/3β5β11βOH-THP, 3α5β11oxo-THP, and 3α5α/3β5β11oxo-THP. Samples were collected in the third trimester, at delivery, and 11-18 months postpartum and quantified by UHPLC-MS ($n = 26$ fecal samples per time point). Concentrations are reported as fmol/mg wet mass (approximately nM). Bars represent mean ± SEM.

(B) Fecal concentrations of progesterone, estradiol, and estrone in the same longitudinal pregnancy cohort ($n = 26$ per time point). Steroids were quantified by UHPLC-MS. Bars represent mean ± SEM.

One-way repeated measures ANOVA with Geisser-Greenhouse correction and Tukey's multiple-comparison test was used for all multi-group comparisons.

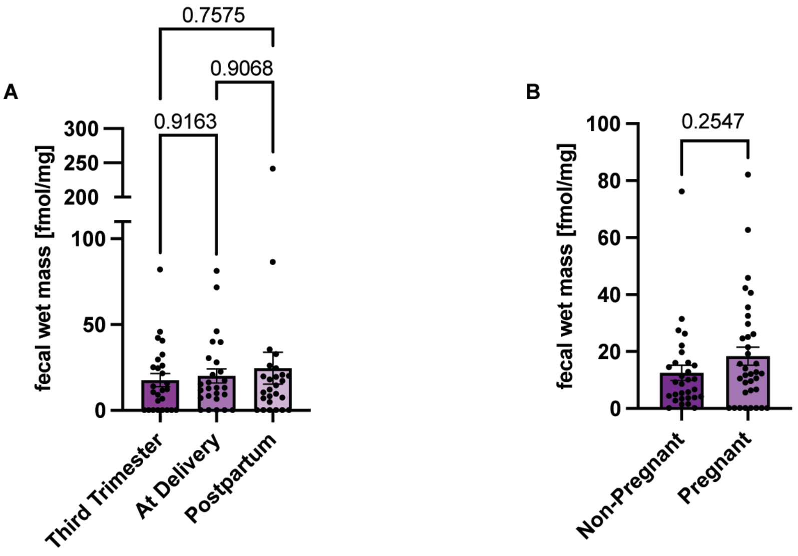


**Supplementary Figure S8. Additional cohort-level comparisons of GALF metabolite 2.**

(A) Fecal levels of **2** across third trimester, delivery, and postpartum time points in the Finnish pregnancy cohort (n = 26 per time point). One-way repeated measures ANOVA with Geisser-Greenhouse correction and Tukey's multiple-comparison test.

(B) Comparison of fecal levels of **2** between pregnant individuals in the Finnish cohort (third trimester from samples with longitudinal data (n=26) with additional third trimester samples without longitudinal data (n=9); total (n = 35)) and female non-pregnant Finnish controls (n = 30). Unpaired two-tailed t test.

All metabolites were quantified by UHPLC-MS and are reported as fmol/mg wet mass. Bars represent mean ± SEM; p values are shown on the plots.

| | Gene Name | | | | |
|---|---|---|---|---|---|
| **Strain Name** | **Elen_1987** | **Elen_2188** | **Elen_1208** | **Elen_1205** | **Elen_2515** |
| **11C** | **0.0** | **0.0** | **0.0** | **0.0** | **0.0** |
| **14A** | **0.0** | **0.0** | **0.0** | **0.0** | **0.0** |
| **16A** | **0.0** | **0.0** | **0.0** | **0.0** | **0.0** |
| **22C** | **0.0** | **0.0** | **0.001** | **0.003** | **0.0** |
| **11863** | **0.0** | **0.0** | **0.001** | **0.003** | **0.0** |
| **25559** | **0.0** | **0.0** | **0.0** | **0.0** | **0.0** |
| **32-6-1-6** | **0.0** | **0.0** | **0.0** | **0.0** | **0.0** |
| **AB12 #2** | **0.0** | **0.0** | **0.0** | **0.0** | **0.0** |
| **CC7/5** | **0.0** | **0.0** | **0.0** | **0.0** | **0.0** |
| **CC8/2** | **0.0** | **0.0** | **2e-6** | **0.003** | **0.0** |
| **CC8/6** | **0.0** | **0.0** | **0.001** | **0.003** | **0.0** |
| **DSM 2243** | **0.0** | **0.0** | **0.0** | **0.0** | **0.0** |
| **UCSF 2243** | **0.0** | **0.0** | **0.0** | **0.0** | **0.0** |
| **Valencia** | **0.0** | **0.0** | **0.001** | **0.003** | **0.0** |
| **W1 BHI6** | **0.0** | **0.0** | **0.001** | **0.003** | **0.0** |

**Supplementary Table 1. Distribution of candidate 11β-hydroxysteroid dehydrogenase genes among producer *Eggerthella lenta* strains**

Presence of putative hydroxysteroid dehydrogenase (HSDH) genes in *E. lenta* strains possessing 11β-hydroxysteroid dehydrogenase (11βHSDH) activity as defined by over 2μM conversion of **1** to **2**. Gene candidates were identified by sequence homology to characterized HSDHs from *E.*

*lenta* DSM 2243 (**see Fig. 3E**). Columns list BLASTn-based E-value scores for each candidate gene (Elen_1987, Elen_2188, Elen_1208, Elen_1205, Elen_2515) across individual strains. Higher values indicate lower nucleotide similarity to the indicated DSM 2243 gene; 0.0 indicates near 100% homolog under the search criteria. Homologs for Elen_1987, Elen_2188, and Elen_2515 were present in all producer strains, and these genes were therefore selected for heterologous expression.
