## Supplemental Data Tables for "Human Gut Bacteria Convert Endogenous Steroids into Host Cortisol Shuttle Inhibitors"

**Supplemental Data S1. Bacterial, cell-assay, human, and mouse data. Related to Figures 1, 2, 4, 5, S1, S3, S5-8, and Star Methods**

Data Table 1: Analyte-specific parameters used in substrate measurement for cell-based assays by UPLC-MS/MS. Related to Fig 2, Fig 4, Fig S3, S5, S6 and STAR Methods.

Data Table 2. Engraftment efficiency of *E. lenta* from FMT of feces from pregnant and nonpregnant human donors into GF mice at day 28. (Adapted from McCurry *et al.* 2024)[49]. Related to Figure S3.

Data Table 3. Primer Sequences for organoid and SW-620 cell-based assays used in RT-qPCR experiments. Related to Fig 4, Fig S5 and STAR Methods.

Data Table 4: Metadata for healthy human donors from a Boston cohort. Related to Fig 1, Fig 5, Fig S1, Fig S8 and STAR methods.

Data Table 5: Metadata for pregnant and non-pregnant human donors from Finland. Related to Fig 5, Fig S7, Fig S8 and STAR methods.

Data Table 6. Metadata for patients with a history of colorectal adenomas from Massachusetts General Hospital. Related to Fig 5 and STAR methods

Data Table 1: Analyte-specific parameters used in substrate measurement for cell-based assays by UPLC-MS/MS. Related to Fig 2, Fig 4, Fig S3, S5, S6 and STAR Methods.

| **Compound** | **Retention times (RT)** | **Precursor ions (Q1)** | **Quantifier-transition** | **Collision energy (CE)** | **qualifier-transition** | **Collision energy (CE)** |
|---|---|---|---|---|---|---|
| Cortisone | 3.28 min | m/z 361.2 | m/z 121.1 | 30 V | m/z 163.1 | 30 V |
| Cortisol | 3.8 min | m/z 363.2 | m/z 97.2 | 44 V | m/z 121 | 30 V |
| Cortisol-D4 | 3.81 min | m/z 367.2 | m/z 120.9 | 36 V | m/z 96.9 | 36 V |
| **2** | 6.27 min | m/z 333.2 | m/z 315.1 | 15 V | m/z 297.2 | 15 V |
| 3α5α11βOH-THP | 8.15 min | m/z 335.2 | m/z 299.1 | 15 V | m/z 105 | 60 V |
| **1** | 8.44 min | m/z 335.3 | m/z 84.9 | 21 V | m/z 91.1 | 60 V |

Data Table 2. Engraftment efficiency of *E. lenta* from FMT from pregnant and nonpregnant human donors into GF mice at day 28. (Adapted from McCurry *et al.* 2024)[49]. Related to Figure S3.

| **Experimental Group** | **Primer pair** | **Avg Cq Value** |
|---|---|---|
| PBS-gavaged | *E. lenta* 16S | 35.79 ± 0.75 |
| Pregnant donor FMT | *E. lenta* 16S | 24.73 ± 0.62 |
| Non-pregnant donor FMT | *E. lenta* 16S | 24.51 ± 0.61 |

Data Table 3. Primer Sequences for organoid and SW-620 cell-based assays used in RT-qPCR experiments. Related to Fig 4, Fig S5 and STAR Methods.

| **Gene** | **Forward Primer 5’-3’** | **Reverse Primer 5’-3’** |
|---|---|---|
| *PPIA* | CATCTGCACTGCCAAGACTGA | TGCAATCCAGCTAGGCATG |
| *HSD11B1* | TCTCAACCACATCACCAACAC | CAGCAACCATTGGATAAGCC |
| *HSD11B2* | CTTCAAGACAGAGTCAGTGAG | GTAGTAGTGGATGAAGTACATGAG |
| *H6PD* | CTGTCCGATTACTACGCCTA | TGAACTCAAAGTCCAACA GA |
| GR (*NR3C1*) | ACAGCATCCCTTTCTCAACAG | AGCTTACATCTGGTCTCATGC |
| MR (*NR3C2*) | AGT GGA AGG GCA ACA CAA CT | TTC TTT GAC TTT CGT GCT CCT |

Data Table 4: Metadata for healthy human donors from a Boston cohort. Related to Fig 1, Fig S1 and STAR methods.

| Patient ID | Sex | Age | Race | Diet | Replicate |
|---|---|---|---|---|---|
| 1 | M | 31 | caucasian | omnivore | A |
| 3 | F | 30 | caucasian | omnivore | |
| 4 | F | 30 | asian | omnivore | N |
| 5 | M | 32 | caucasian | omnivore | A |
| 6 | M | 27 | asian | omnivore | I |
| 7 | M | 30 | caucasian-african | omnivore | EE |
| 8 | F | 26 | caucasian | omnivore | F |
| A | M | 31 | caucasian | omnivore | |
| AA | M | 25 | asian | omnivore | |
| BB | F | 24 | caucasian-asian | vegetarian | |
| C | F | 28 | caucasian | omnivore | |
| CC | F | 26 | caucasian | omnivore | |
| D | M | 30 | caucasian | omnivore | |
| DD | F | 27 | caucasian | omnivore; no seafood, fish or pork | |
| EE | M | 29 | caucasian-african | omnivore | |
| F | F | 24 | caucasian | omnivore | |
| FF | M | 28 | asian | omnivore | |
| G | M | 28 | caucasian | omnivore | |
| GG | M | 30 | caucasian | omnivore | |
| H | M | 26 | caucasian | vegan | |
| I | M | 26 | asian | omnivore | |
| II | M | 38 | middle east | omnivore | |
| J | F | 27 | caucasian | omnivore | |
| K | M | 27 | caucasian | omnivore | |
| L | F | 23 | asian | omnivore | |
| M | F | 28 | caucasian | omnivore | |
| N | F | 29 | asian | omnivore | |
| O | M | 32 | caucasian | omnivore | |
| P | M | 24 | caucasian | omnivore | |
| Q | M | 28 | caucasian | omnivore | |
| R | F | 27 | caucasian | pescetarian | |

| | | | | |
|---|---|---|---|---|
| S | M | 28 | indian | omnivore |
| T | M | 31 | caucasian | omnivore |
| U | F | 32 | caucasian | omnivore, gluten-free |
| V | F | 29 | caucasian | omnivore |
| W | F | 34 | caucasian | omnivore, medicated |
| X | M | 27 | caucasian | omnivore |
| Y | M | 26 | caucasian | omnivore |
| Z1 | M | 27 | caucasian | omnivore |

Data Table 5: Metadata for Finland Human Cohort Donors. Related to Fig 5, Fig S7, Fig S8, and STAR methods.

| **Participant** | **Sample ID** | **Age** | **Pregnancy Status** | **Longitudinal** | **Chronic disease** | **disease diagnosed during pregnancy/abx or glucocorticoids during pregnancy** |
|---|---|---|---|---|---|---|
| M0018M | 300034 | 32.5 | 0 months | yes | | |
| M0018M | 900084 | 33.5 | 12 months | yes | | |
| M0018M | 300004 | 32.3 | During pregnancy | yes | | GDM, insulin treatment |
| M0024M | 300046 | 34.5 | 0 months | yes | | |
| M0024M | 900145 | 35.7 | 14 months | yes | | |
| M0024M | 300013 | 34.2 | During pregnancy | yes | Celiac disease | |
| M0333M | 300225 | 31.5 | During pregnancy | yes | Migraine | |
| M0333M | 302748 | 33.3 | 18 months | yes | | |
| M0333M | 300365 | 31.8 | 0 months | yes | | |
| M0346M | 300239 | 29.3 | During pregnancy | yes | | |
| M0346M | 300390 | 29.5 | 0 months | yes | | |
| M0346M | 302756 | 31.1 | 18 months | yes | | |

| | | | | | | |
|---|---|---|---|---|---|---|
| M0353M | 302621 | 29.5 | 18 months | yes | | |
| M0353M | 301046 | 27.4 | 0 months | yes | | |
| M0353M | 300238 | 27.2 | During pregnancy | yes | | |
| M0388M | 302793 | 29.6 | 18 months | yes | | |
| M0388M | 300420 | 28.1 | 0 months | yes | | |
| M0388M | 300277 | 27.8 | During pregnancy | yes | | |
| M0450M | 302717 | 38.7 | 18 months | yes | | |
| M0450M | 301266 | 37.2 | 0 months | yes | | |
| M0450M | 300429 | 37.0 | During pregnancy | yes | | GDM |
| M0552M | 300515 | 31.1 | 0 months | yes | | |
| M0552M | 300362 | 30.9 | During pregnancy | yes | | |
| M0552M | 302711 | 32.7 | 18 months | yes | | |
| M0740M | 301404 | 35.8 | 0 months | yes | | |
| M0740M | 302743 | 37.3 | 18 months | yes | | |
| M0740M | 300494 | 35.6 | During pregnancy | yes | | |
| M0744M | 301563 | 32.3 | 0 months | yes | | |
| M0744M | 301105 | 32.0 | During pregnancy | yes | Acne | |
| M0744M | 302633 | 33.3 | 12 months | yes | | |
| M0757M | 302637 | 46.8 | 12 months | yes | | |
| M0757M | 301059 | 45.6 | During pregnancy | yes | | |
| M0757M | 301290 | 45.8 | 0 months | yes | | |
| M0806M | 301062 | 33.4 | During pregnancy | yes | | |
| M0806M | 302758 | 34.7 | 12 months | yes | | |

| | | | | | | |
|---|---|---|---|---|---|---|
| M0806M | 301260 | 33.6 | 0 months | yes | | |
| M0808M | 301223 | 33.8 | During pregnancy | yes | | |
| M0808M | 301432 | 34.0 | 0 months | yes | | |
| M0808M | 302664 | 35.0 | 12 months | yes | | |
| M0893M | 301293 | 26.9 | 0 months | yes | | |
| M0893M | 301122 | 26.7 | During pregnancy | yes | | |
| M0893M | 302776 | 27.9 | 12 months | yes | | |
| M0969M | 302623 | 39.3 | 12 months | yes | | |
| M0969M | 301308 | 38.4 | 0 months | yes | | |
| M0969M | 301094 | 38.1 | During pregnancy | yes | | |
| M1066M | 302672 | 25.6 | 12 months | yes | | |
| M1066M | 301721 | 24.6 | 0 months | yes | | |
| M1066M | 301227 | 24.4 | During pregnancy | yes | Hypothyroidism | |
| M1097M | 302742 | 36.4 | 12 months | yes | | |
| M1097M | 301399 | 35.1 | During pregnancy | yes | | GDM |
| M1097M | 301884 | 35.3 | 0 months | yes | | |
| M1172M | 301643 | 42.3 | 0 months | yes | | |
| M1172M | 302803 | 43.5 | 12 months | yes | | |
| M1172M | 301465 | 42.2 | During pregnancy | yes | | |
| M1181M | 302741 | 45.3 | 12 months | yes | | |
| M1181M | 301474 | 44.2 | During pregnancy | yes | | |
| M1181M | 301703 | 44.4 | 0 months | yes | | |
| M1294M | 302770 | 32.0 | 12 months | yes | | |

| | | | | | | |
|---|---|---|---|---|---|---|
| M1294M | 301770 | 31.0 | 0 months | yes | | |
| M1294M | 301528 | 30.8 | During pregnancy | yes | | |
| M1300M | 301539 | 25.9 | During pregnancy | yes | | |
| M1300M | 302781 | 27.2 | 12 months | yes | | |
| M1300M | 301786 | 26.1 | 0 months | yes | | |
| M1301M | 301463 | 28.0 | During pregnancy | yes | Colon irritable, no medication | |
| M1301M | 302760 | 29.2 | 12 months | yes | | |
| M1301M | 301950 | 28.2 | 0 months | yes | | |
| M1332M | 301801 | 29.9 | 0 months | yes | | |
| M1332M | 302797 | 30.9 | 12 months | yes | | |
| M1332M | 301595 | 29.7 | During pregnancy | yes | Gastroesophageal reflux disease | |
| M1381M | 301720 | 25.0 | During pregnancy | yes | | UTI, antibiotics |
| M1381M | 302799 | 26.2 | 12 months | yes | | |
| M1381M | 301899 | 25.2 | 0 months | yes | | |
| M1409M | 302727 | 37.7 | 12 months | yes | | |
| M1409M | 301464 | 36.6 | During pregnancy | yes | | |
| M1409M | 301682 | 36.7 | 0 months | yes | | |
| M1443M | 301611 | 38.7 | During pregnancy | yes | | |
| M1443M | 302785 | 39.8 | 12 months | yes | | |
| M1443M | 301981 | 38.9 | 0 months | yes | | |
| M0041M | 300011 | 31.7 | During pregnancy | no | | |
| M0083M | 300028 | 29.1 | During pregnancy | no | | |
| M0092M | 300035 | 30.5 | During pregnancy | no | | |

| | | | | |
|---|---|---|---|---|
| M0107M | 300040 | 31.2 | During pregnancy | no |
| M0115M | 300050 | 34.9 | During pregnancy | no |
| M0121M | 300056 | 31.5 | During pregnancy | no |
| M0126M | 300061 | 37.9 | During pregnancy | no |
| M0139M | 300068 | 28.4 | During pregnancy | no |
| M0149M | 300064 | 40.1 | During pregnancy | no |
| M0001E | 900046 | 31.0 | Non-pregnant | no |
| M0006E | 900047 | 35.6 | Non-pregnant | no |
| M0002E | 900056 | 27.2 | Non-pregnant | no |
| M0011E | 900057 | 29.8 | Non-pregnant | no |
| M0012E | 900058 | 24.9 | Non-pregnant | no |
| M0009E | 900061 | 36.6 | Non-pregnant | no |
| M0003E | 900062 | 25.5 | Non-pregnant | no |
| M0014E | 900069 | 28.2 | Non-pregnant | no |
| M0008E | 900070 | 27.2 | Non-pregnant | no |
| M0017E | 900072 | 33.8 | Non-pregnant | no |
| M0007E | 900077 | 25.0 | Non-pregnant | no |
| M0016E | 900078 | 40.3 | Non-pregnant | no |
| M0018E | 900084 | 36.2 | Non-pregnant | no |
| M0019E | 900085 | 41.9 | Non-pregnant | no |
| M0005E | 900086 | 32.3 | Non-pregnant | no |
| M0004E | 900091 | 32.3 | Non-pregnant | no |
| M0021E | 900099 | 34.4 | Non-pregnant | no |

| | | | | |
|---|---|---|---|---|
| M0022E | 900102 | 38.3 | Non-pregnant | no |
| M0023E | 900136 | 38.1 | Non-pregnant | no |
| M0024E | 900145 | 35.4 | Non-pregnant | no |
| M0027E | 900161 | 30.3 | Non-pregnant | no |
| M0010E | 900212 | 35.0 | Non-pregnant | no |
| M0029E | 900223 | 23.7 | Non-pregnant | no |
| M0028E | 900227 | 41.9 | Non-pregnant | no |
| M0026E | 900229 | 29.5 | Non-pregnant | no |
| M0025E | 900246 | 23.5 | Non-pregnant | no |
| M0030E | 900269 | 26.4 | Non-pregnant | no |
| M0031E | 900279 | 32.4 | Non-pregnant | no |
| M0032E | 900282 | 27.9 | Non-pregnant | no |
| M0033E | 900283 | 29.0 | Non-pregnant | no |

GDM = Gestational Diabetes; UTI = urinary tract infection

Data Table 6. Metadata for patients with colorectal adenomas at Massachusetts General Hospital. Related to Fig 5 and STAR methods.

| Subject | Age | Gender | Systolic | Diastolic | Hypertension-range (not diagnostic) | BMI | Antihypertensive medication |
|---|---|---|---|---|---|---|---|
| 2 | 78 | M | 147 | 78 | stage-2 | 30 | yes |
| 4 | 75 | M | 167 | 81 | stage-2 | 32 | |
| 5 | 76 | M | 167 | 99 | stage-2 | 25 | no |
| 7 | 66 | M | 129 | 58 | elevated | 35 | yes |
| 12 | 67 | M | 122 | 79 | elevated | 21 | |
| 16 | 74 | F | 121 | 74 | elevated | 18 | no |

| | | | | | | | |
|---|---|---|---|---|---|---|---|
| 19 | 75 | M | 148 | 72 | stage-2 | 27 | |
| 20 | 78 | M | 126 | 67 | elevated | 28 | yes |
| 22 | 74 | F | 154 | 75 | stage-2 | 38 | |
| 25 | 74 | F | 137 | 87 | stage-1 | 26 | yes |
| 30 | 62 | M | 148 | 72 | stage-2 | 25 | no |
| 33 | 76 | M | 152 | 76 | stage-2 | 28 | yes |
| 40 | 72 | M | 149 | 92 | stage-2 | 26 | yes |
| 49 | 72 | M | 144 | 90 | stage-2 | 24 | no |
| 51 | 55 | M | 133 | 79 | stage-1 | 32 | |
| 52 | 60 | M | 132 | 78 | stage-1 | 26 | |
| 58 | 66 | F | 149 | 67 | stage-2 | 26 | yes |
| 59 | 67 | F | 150 | 76 | stage-2 | 23 | no |
| 60 | 58 | M | 140 | 85 | stage-2 | 35 | no |
| 64 | 49 | M | 122 | 63 | elevated | 22 | |
| 66 | 70 | M | 160 | 75 | stage-2 | 40 | yes |
| 67 | 62 | M | 135 | 95 | stage-2 | 35 | no |
| 68 | 65 | M | 168 | 118 | stage-2 | 30 | no |
| 70 | 58 | M | 147 | 88 | stage-2 | 25 | |
| 71 | 76 | M | 146 | 76 | stage-2 | 26 | no |
| 77 | 45 | M | 122 | 74 | elevated | 23 | |
| 78 | 58 | F | 119 | 70 | normal | 23 | |
| 79 | 56 | M | 118 | 74 | normal | 28 | no |
| 80 | 50 | F | 124 | 56 | elevated | 22 | |
| 81 | 60 | M | 150 | 85 | stage-2 | 30 | yes |
| 86 | 75 | F | 179 | 76 | stage-2 | 32 | no |

| | | | | | | | |
|---|---|---|---|---|---|---|---|
| 88 | 64 | M | 138 | 88 | stage-1 | 23 | yes |
| 89 | 63 | M | 156 | 88 | stage-2 | 26 | yes |
| 90 | 51 | F | 123 | 81 | stage-1 | 28 | no |
| 92 | 54 | M | 121 | 84 | stage-1 | 23 | |
| 93 | 69 | F | 153 | 76 | stage-2 | 30 | yes |
| 94 | 47 | M | 135 | 79 | stage-1 | 27 | |
| 95 | 62 | F | 143 | 84 | stage-2 | 34 | |
| 97 | 67 | M | 144 | 82 | stage-2 | 29 | |
| 98 | 65 | M | 152 | 75 | stage-2 | 26 | |
| 100 | 76 | F | 156 | 105 | stage-2 | 39 | yes |
| 103 | 71 | F | 124 | 60 | elevated | 33 | |
| 104 | 53 | M | 132 | 87 | stage-1 | 29 | |
| 105 | 53 | M | 124 | 81 | stage-1 | 33 | |
| 108 | 71 | M | 147 | 76 | stage-2 | 25 | no |
| 109 | 58 | M | 169 | 85 | stage-2 | 30 | no |
| 111 | 49 | M | 141 | 87 | stage-2 | 29 | |
| 113 | 65 | M | 182 | 88 | stage-2 | 35 | no |
| 114 | 55 | M | 115 | 71 | normal | 26 | |
| 116 | 63 | M | 139 | 79 | stage-1 | 32 | yes |
| 117 | 61 | M | 134 | 65 | stage-1 | 27 | yes |
| 119 | 56 | F | 128 | 83 | stage-1 | 33 | |
| 122 | 50 | F | 138 | 89 | stage-1 | 22 | |
